# Steroid hormone signaling activates thermal nociception during *Drosophila* peripheral nervous system development

**DOI:** 10.1101/2021.12.02.470982

**Authors:** Jacob S. Jaszczak, Laura DeVault, Lily Yeh Jan, Yuh Nung Jan

## Abstract

Sensory neurons enable animals to detect environmental changes and avoid harm. An intriguing open question concerns how the various attributes of sensory neurons arise in development. *Drosophila melanogaster* larvae undergo a behavioral transition by robustly activating a thermal nociceptive escape behavior during the second half of larval development (3^rd^ instar). The Class 4 dendritic arborization (C4da) neurons are multimodal sensors which tile the body wall of *Drosophila* larvae and detect nociceptive temperature, light, and mechanical force. In contrast to the increase in nociceptive behavior in the 3^rd^ instar, we find that ultraviolet light-induced Ca^2+^ activity in C4da neurons decreases during same period of larval development. Loss of ecdysone receptor has previously been shown to reduce nociception in 3^rd^ instar larvae. We find that ligand dependent activation of ecdysone signaling is sufficient to promote nociceptive responses in 2^nd^ instar larvae and suppress expression of *subdued* (encoding a TMEM16 channel). Reduction of *subdued* expression in 2^nd^ instar C4da neurons not only increases thermal nociception but also decreases the response to ultraviolet light. Thus, steroid hormone signaling suppresses *subdued* expression to facilitate the sensory switch of C4da neurons. This regulation of a developmental sensory switch through steroid hormone regulation of channel expression raises the possibility that ion channel homeostasis is a key target for tuning the development of sensory modalities.

## Introduction

Detection of internal and external stimuli depends on developmental programing of the proper physiological and morphological properties of sensory neurons. It is an intriguing open question as to how the characteristics of sensory transduction arise during development, and which neuronal properties are crucial for the initiation of sensory detection.

Nociception, the ability to detect and escape potentially harmful stimuli, is a highly conserved function of sensory systems present in all animals (Arenas et al. 2017). A nociception system is composed of nociceptors, which are neurons with molecular components that directly sense harmful stimuli; a processing center where the stimuli are interpreted, typically the neural circuits of the central nervous system (CNS); and the reflexive neuromuscular circuit which generates a reaction to escape injury (Baliki and Apkarian 2015). Development of sensory systems requires the coordination of processes on the cellular, morphogenic, and circuit level.

Thermal nociception generates an avoidance behavior, such as a fast limb retraction or change of movement trajectory. Previous studies of thermal nociception behaviors and neural activity demonstrate a change in nociceptive responses during development of both mice and *Drosophila*. For example, a mouse paw exhibits low sensitivity to painful heat early in postnatal development, but the rate of reflex gets faster as the animal develops (Jankowski et al. 2014; Ford et al. 2019). A developmental transition is also observed in nociceptor activity, as revealed by the change in the number of heat sensitive C-Fibers with age in mice (Hiura et al. 1992). Likewise, a behavioral transition in heat thermal nociception has been observed in *Drosophila melanogaster* larvae (Sulkowski et al. 2011). *Drosophila* 3^rd^ instar larvae exhibit a stereotyped “corkscrew” behavior, when a thermal probe heated above 38°C touches the cuticle (Tracey et al. 2003; Babcock et al. 2009). However, this behavior is not observed in 2^nd^ instar or earlier larval stages, and appears to be acquired only during the last stage of larval development.

The larval nociceptive behavior has been a useful model for dissection of the molecular mechanisms and circuitry of nociception (Himmel et al. 2017). Among the heat sensitive neurons in the larval cuticle (Liu et al., 2003) are the Class IV dendritic arborization (C4da) neurons, which function as the primary nociceptors for noxious heat induced escape behavior (Tracey et al. 2003; Hwang et al. 2007). C4da neurons are multimodal sensors for harmful stimuli, with the capacity to respond to multiple forms of sensory information. In addition to responding to temperature, C4da neurons respond to high intensity blue light (Xiang et al. 2010), noxious UV light (Xiang et al. 2010; Gu et al. 2019), and harsh touch (Zhong et al. 2010).

While noxious heat and touch both elicit the rolling escape behavior, stimulation of C4da neurons with short wavelength (blue or UV) light causes a locomotion trajectory avoidance behavior. These different stimuli cause different electrical signals in C4da neurons. Noxious heat causes high-frequency spike trains interspersed with quiescent periods, in contrast to blue light which elicits lower frequency firing with few quiescent periods in 3^rd^ instar larvae (Terada et al. 2016). Therefore, distinct behaviors appear to be encoded by the C4da neurons according to their differential responses to differing stimuli.

The majority of studies of nociception in *Drosophila* larvae have focused on the last (i.e., the 3^rd^) larval instar, where nearly all larvae will perform the behavioral response. However, Sulkowski and colleagues found that during the first half of larval development (i.e., the 1^st^ and the 2^nd^ instar), larvae exhibited either no or a low rate of thermal nociceptive behavior, leading to the hypothesis that a developmental transition occurs between the 2^nd^ and the 3^rd^ larval instar (Sulkowski et al. 2011). The mechanisms which regulate this transition remain unknown. Optogenetic activation of C4da neurons can induce the corkscrew behavior or a locomotion trajectory avoidance behavior in 3^rd^ instar larvae. However, optogenetic activation could only induce a locomotion trajectory avoidance behavior in the 2^nd^ instar. This difference between 2^nd^ and 3^rd^ instar in the optogenetic activation of behavior, taken together with the difference in coding activity in C4da neurons in response to thermal stimulation or light (Terada et al. 2016), raises the possibility that the developmental transition may involve a change in sensory coding. Supporting this possibility, misexpression of low temperature activated TrpA1 (isoform A) channels in the C4da neurons enables 2^nd^ instar larvae to exhibit corkscrew behavior in response to an innocuous temperature stimulus (Luo et al. 2017). It is conceivable that the sensory switch may be caused by a cell autonomous change in the C4da neurons. Whether C4da neurons change in their neural activity during the developmental transition from 2^nd^ to 3^rd^ instar is unknown, and the regulators of such transition are unknown as well.

Transcription factors facilitate the formation of distinct neuronal functions (Parrish et al. 2014) and regulate the dynamic expression profiles of many developmental transitions between and during larval instars (Sullivan and Thummel 2003). One such transcription factor is the nuclear hormone receptor Ecdysone Receptor (EcR). EcR forms a heterodimer with Ultraspiracle (USP) and binds the ligand ecdysone, a steroid hormone, to coordinate a wide variety of developmental programs (Yamanaka et al. 2013). EcR and USP are required for thermal nociception in the 3^rd^ instar (McParland et al. 2015), but the mechanism and role of ecdysone signaling in the 2^nd^ instar has not been determined.

EcR expression in C4da neurons has been reported at the beginning and end of larval development (Kuo et al. 2005; Ou et al. 2008; Kirilly et al. 2009; McParland et al. 2015). In the absence of ligand, EcR and USP distribute between the cytoplasm and nucleus, while ligand binding increases nuclear localization (Nieva et al. 2007, 2008). Without ligand binding, EcR and USP recruit nucleosome remodeling proteins to compact the chromatin and locally suppress transcription. When bound to ecdysone, EcR-USP recruit coactivators, such as histone acetylases, to expand the chromatin structure and promote transcription. In this way steroid hormone signaling can create positive and negative feedback to promote cascades of transcriptional programs at developmentally precise periods (Sullivan and Thummel 2003; Hill et al. 2013). Whether the ligand-induced activity and transcriptional regulation are involved in EcR mediated thermal nociception remains to be determined.

In this study, we performed a series of experiments to determine the activity of the C4da neurons during the developmental nociceptive transition as well as the mechanisms which regulate this sensory switch. These experiments demonstrate that steroid hormone signaling in nociceptive neurons regulates the channel composition in C4da neurons to modulate the sensory properties and behavioral response at distinct stages of larval development.

## Results

### C4da neuron sensory switch between 2^nd^ and 3^rd^ larval instar is modality specific

C4da neurons are multimodal nociceptive sensors of thermal, short wavelength light, and mechanical stimuli (Himmel et al. 2017). While 1^st^ instar and 3^rd^ instar larvae respond similarly to mechano-nociceptive stimuli (Almeida-Carvalho et al. 2017), larvae do not robustly activate the thermally triggered nociceptive behavior until the last (3^rd^) instar of development (Sulkowski et al. 2011). Whether the C4da neurons respond to short wavelength light changes during larval development is unknown. To investigate whether this developmental transition is specific to thermal nociception, we sought to compare the responses of 2^nd^ and 3^rd^ instar larvae to thermal and light stimuli.

First, we measured the frequency and latency of the nociceptive behavior in 2^nd^ and 3^rd^ instar larvae across a range of nociceptive temperatures. Compared to 3^rd^ instar larvae, 2^nd^ instar larvae had a lower response frequency of nociceptive behavior across all nociceptive temperatures above 40°C (Figure 1A). Among the larvae which did display a nociceptive behavior, 2^nd^ instar larvae had a longer latency before initiating the nociceptive response than 3^rd^ instar larvae (Figure 1B, Figure 1 – figure supplement 1A,B). Thus, 2^nd^ instar larvae displayed reduced thermal nociceptive behavior across a wide range of temperatures.

**Figure 1:**
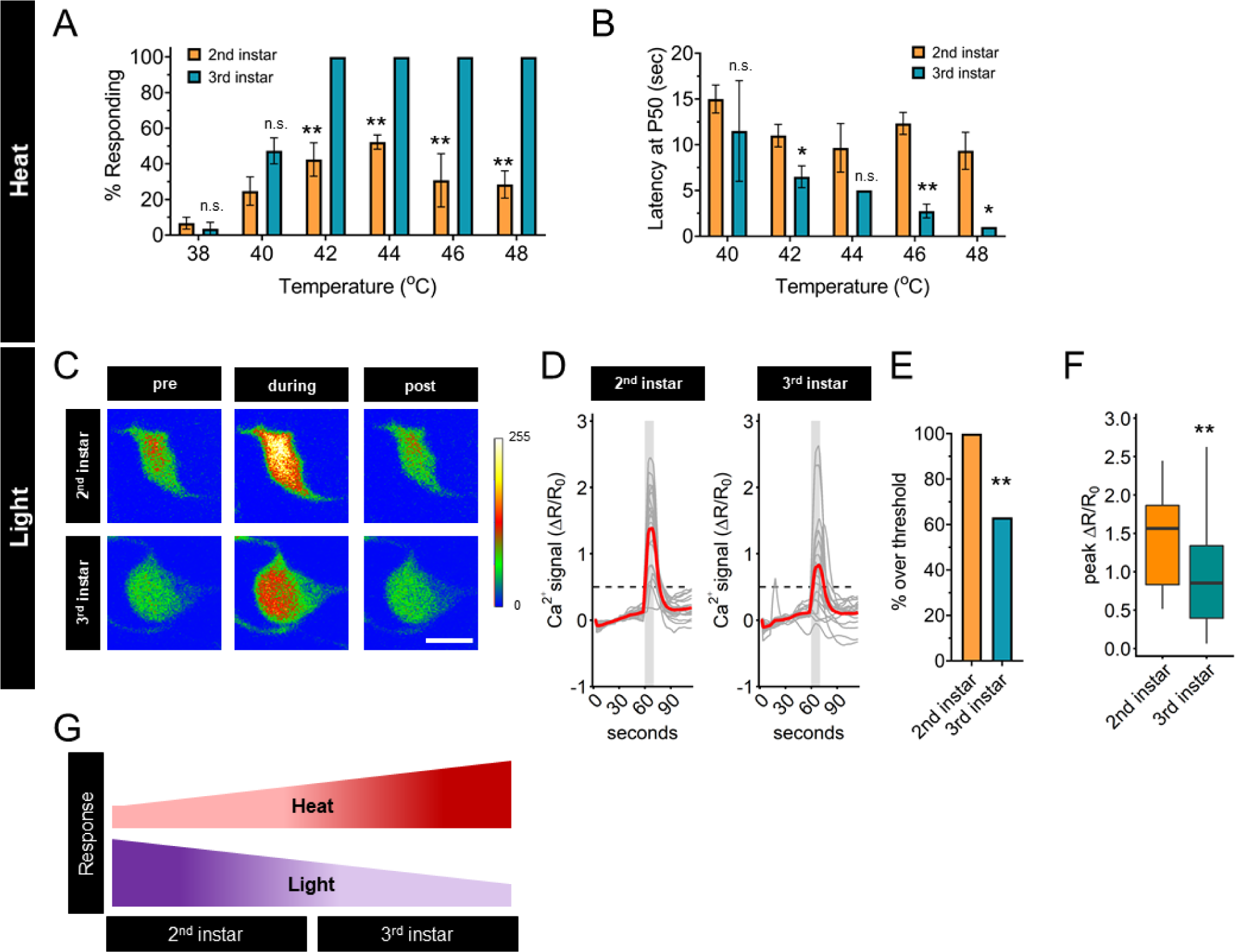
Thermal nociception behaviors increase during larval development, while UV-A light response decreases. (A) Groups of age-matched control larvae were touched with a thermal probe at different temperatures. The nociceptive response was scored if a 360° roll occurred during 20 seconds of stimulus. (B) Latency by which 50% of the responding population of larvae has had a nociceptive behavior response. (C) Representative GCaMP6s images of 2^nd^ and 3^rd^ instar in soma C4da neurons. Scale bar = 10μm. Periods of UV-A treatment program: Pre = before UV-A stimulus. During = period of UV-A light stimulus. Post = after the end of UV-A stimulus. (D) Individual traces (grey lines) and means (red lines) of Ca^2+^ activity calculated by ratiometric change from baseline. Grey column indicates period of UV-A stimulus. Dotted line indicates threshold level for % over threshold calculations. (E) Percent of soma with Ca^2+^ activity over the threshold level. (F) Peak Ca^2+^ activity during the periods of UV-A treatment programs. (G) Heat and light development have opposite trends during larval development. A: n = 2-4 staging replicates of 15-20 larvae were tested for each age and temperature. C-F: 2^nd^ instar n = 21 neurons. 3^rd^ instar n = 19 neurons. A,B: Student’s t-test. E: Fisher’s exact test, one-sided. F: Student’s t-test. * p < 0.05,** p < 0.01. **Figure 1 - figure supplement 1:** Thermal nociceptive sensitivity increases across temperatures from 2^nd^ to 3rd instar.

High intensity UV light exposure of 3^rd^ instar larvae causes behavioral responses, increased Ca^2+^ activity, and patterns of spike trains that are different from those caused by nociceptive temperatures (Xiang et al. 2010; Terada et al. 2016). The small size of 2^nd^ instar larvae makes electrophysiological recording of C4da neurons prohibitive, therefore, we sought to compare the magnitude of the UV-induced Ca^2+^ response. To determine if the UV light-induced Ca^2+^ activity response of C4da neurons changes during development, we measured GCaMP6s signals of these sensory neurons in 2^nd^ and 3^rd^ instar larvae in response to 405 nm (UV-A) light stimulation. In contrast to the transition of thermal nociceptive behavior, UV-A stimuli induced Ca^2+^ activity of C4da neurons in 2^nd^ instar larvae was higher than that in 3rd instar larvae (Figure 1C,D). This pattern, of 3^rd^ instar C4da neurons having lower UV-A induced Ca^2+^ activity than 2^nd^ instar C4da neurons, was validated by quantifying both the number of neurons which surpassed a threshold value of UV-A induced Ca^2+^ activity and the magnitude of the peak Ca^2+^ activity (Figure 1E,F). Thus, the UV-A induced Ca^2+^ activity decreased with larval development, in contrast to the increase of thermal nociceptive neuronal response from 2^nd^ instar to 3^rd^ instar (Figure 1G). Based on these observations, we conclude that regulation of the sensory switch during 2^nd^ and 3^rd^ instar development is modality specific.

### Steroid hormone ecdysone promotes the development of thermal nociception

Having found modality specific sensory responses during the 2^nd^ to 3^rd^ instar developmental transition, we sought to find regulators of the sensory switch in C4da neurons by first examining the thermal nociceptive transition. The steroid hormone ecdysone regulates multiple developmental events during the 2^nd^ and 3^rd^ instar, with peaks in ecdysone titer triggering molting events and developmental checkpoints (Rewitz et al. 2013). Ecdysone is the ligand for the Ecdysone Receptor (EcR) with three isoforms (A, B1, and B2). These isoforms are identical in the DNA binding and ligand binding domains but differ in the Activation Function 1 (AF1) domain (Figure 2 – figure supplement 1A). C4da neurons express isoforms EcR-A and EcR-B1 at embryonic stages, during the 3^rd^ instar, and dynamically during pupal development (Kuo et al. 2005; Ou et al. 2008; Kirilly et al. 2009; McParland et al. 2015). EcR-B1 regulates dendrite growth (Ou et al., 2008) and pruning (Kuo et al., 2005, Kirilly et al., 2009). EcR-A is required for thermal nociception in the 3^rd^ instar, as revealed by the greater reduction of nociception caused by RNAi knockdown of EcR-A as compared to RNAi knockdown of EcR-B1 (McParland et al. 2015). Therefore, we investigated the role of ecdysone and EcR-A in the developmental transition of thermal nociceptive behavior.

**Figure 2:**
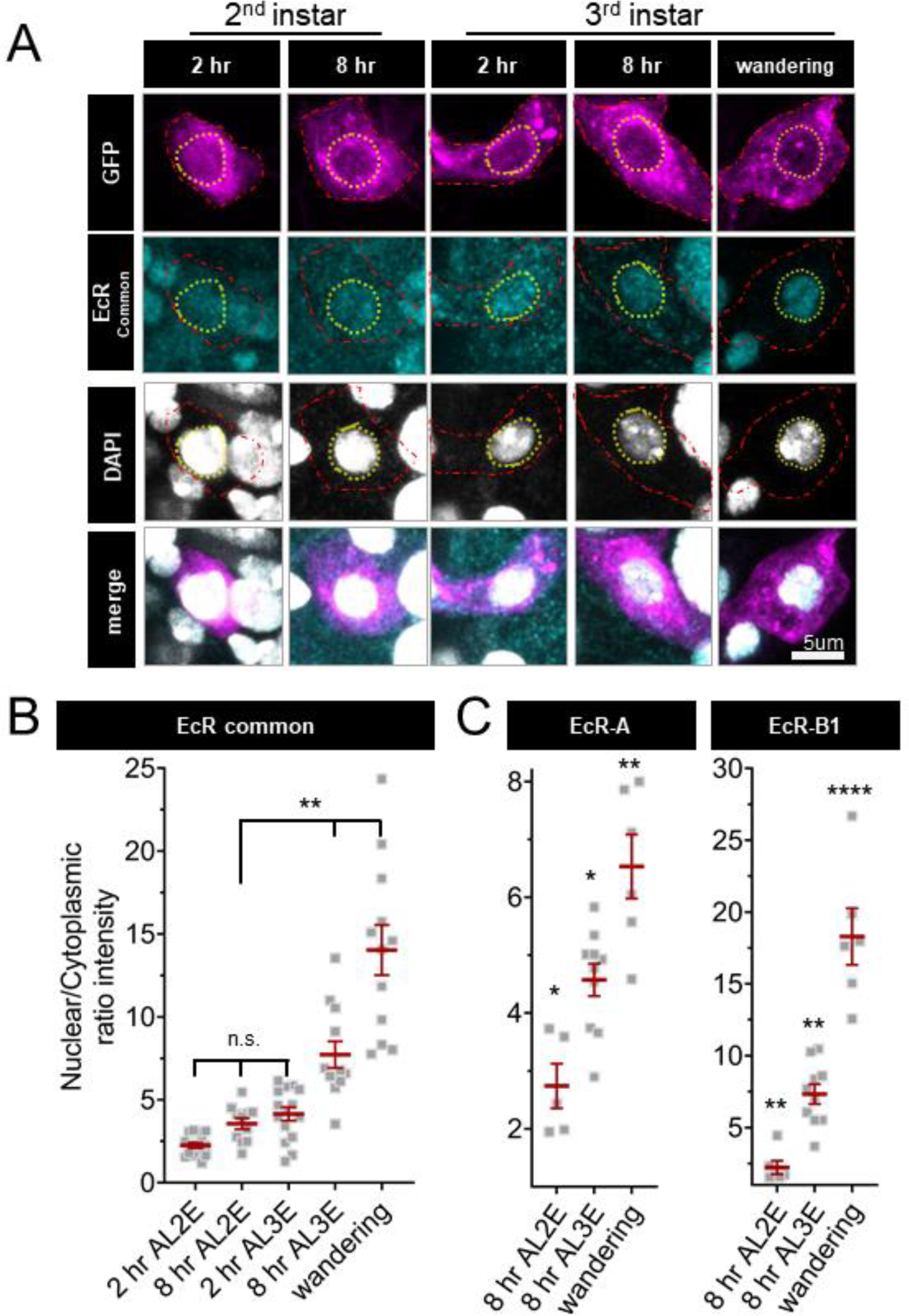
Ecdysone receptor increases nuclear localization in C4da neuron early during the 3^rd^ instar. (A) EcR immunohistochemistry with EcR-common antibody. Red dashed line outlines C4da neurons and yellow dashed line indicates position of nucleus. C4da neurons labeled by ppktd-GFP. EcR nuclear localization reported by nuclear to cytoplasmic ratio of intensity with (B) EcR-common and (C) EcR-A, or EcR-B1 antibody. After L2 Ecdysis (AL2E), After L3 Ecdysis (AL3E). One-way Anova with Bonferroni. * p < 0.05,** p < 0.01,**** p < 0.0001.

Nuclear localization of EcR increases during the end of larval development; localization is greater in late 3^rd^ instar larvae and pupae as compared to early third instar larvae (Kuo et al. 2005; Kirilly et al. 2009). However, the expression and localization of EcR isoforms during the 2^nd^ larval instar is unknown. To investigate EcR dynamics during the developmental period with different levels of thermal nociception, we examined the localization of EcR in 2^nd^ and 3^rd^ instar larval C4da neurons with an antibody which recognizes a common domain of all EcR isoforms. Consistent with previous findings (Kuo et al. 2005; Ou et al. 2008; Kirilly et al. 2009; McParland et al. 2015), we observed nuclear localization of EcR in wandering 3^rd^ instar larvae (Figure 2A,B). Earlier in larval development, EcR was evenly distributed between the nucleus and cytoplasm in the 2^nd^ instar (2 hr and 8 hr After L2 Ecdysis) and at the beginning of the 3^rd^ instar, 2 hr After L3 Ecdysis (AL3E). Nuclear localization increased 8 hrs AL3E and in wandering 3^rd^ instar larvae (Figure 2A,B). Similar EcR localization dynamics could be detected for specific EcR isoforms, as revealed by antibodies specific to EcR-A and EcR-B1, respectively (Figure 2C). Quantification of total EcR immunoreactivity in C4da neurons did not reveal any significant change during 2^nd^ and 3^rd^ instar development. (Figure 2 – figure supplement 1B). These observations suggest that while EcR is present in C4da neurons throughout the 2^nd^ and 3^rd^ instar, the nuclear localization begins to increase early in the 3^rd^ instar.

Changes in ecdysone synthesis and systemic ecdysone titer orchestrate critical periods of *Drosophila* development. Pulses of ecdysone in the 1^st^ and 2^nd^ larval instars increase systemic titers and promote expression of genes required for molting, while a series of three pulses of ecdysone in the 3^rd^ instar control developmental checkpoints and developmental preparation for metamorphosis (Warren et al. 2006; Kannangara et al. 2021). To determine whether the increased thermal nociception in 3^rd^ instar larvae is driven by systemic ecdysone, we tested whether increasing steroid hormone titer during the 2^nd^ instar is sufficient to cause precocious development of thermal nociceptive behavior. Feeding larvae food supplemented with the metabolically active 20-hydroxyecdysone (20E) increases ecdysone signaling in larvae (Colombani et al. 2005). Thus, we transferred larvae to 20E supplemented food and measured larval thermal nociceptive behavior 8 hours later, while the larvae were still in the 2^nd^ instar. Larvae fed with 250 μg/ml 20E had a higher rate of behavioral response to thermal nociceptive stimuli as compared to larvae fed with 125 μg/ml 20E or vehicle alone (Figure 3A), indicating that increasing ecdysone titer in the 2^nd^ instar is sufficient to promote the development of thermal nociception.

**Figure 3:**
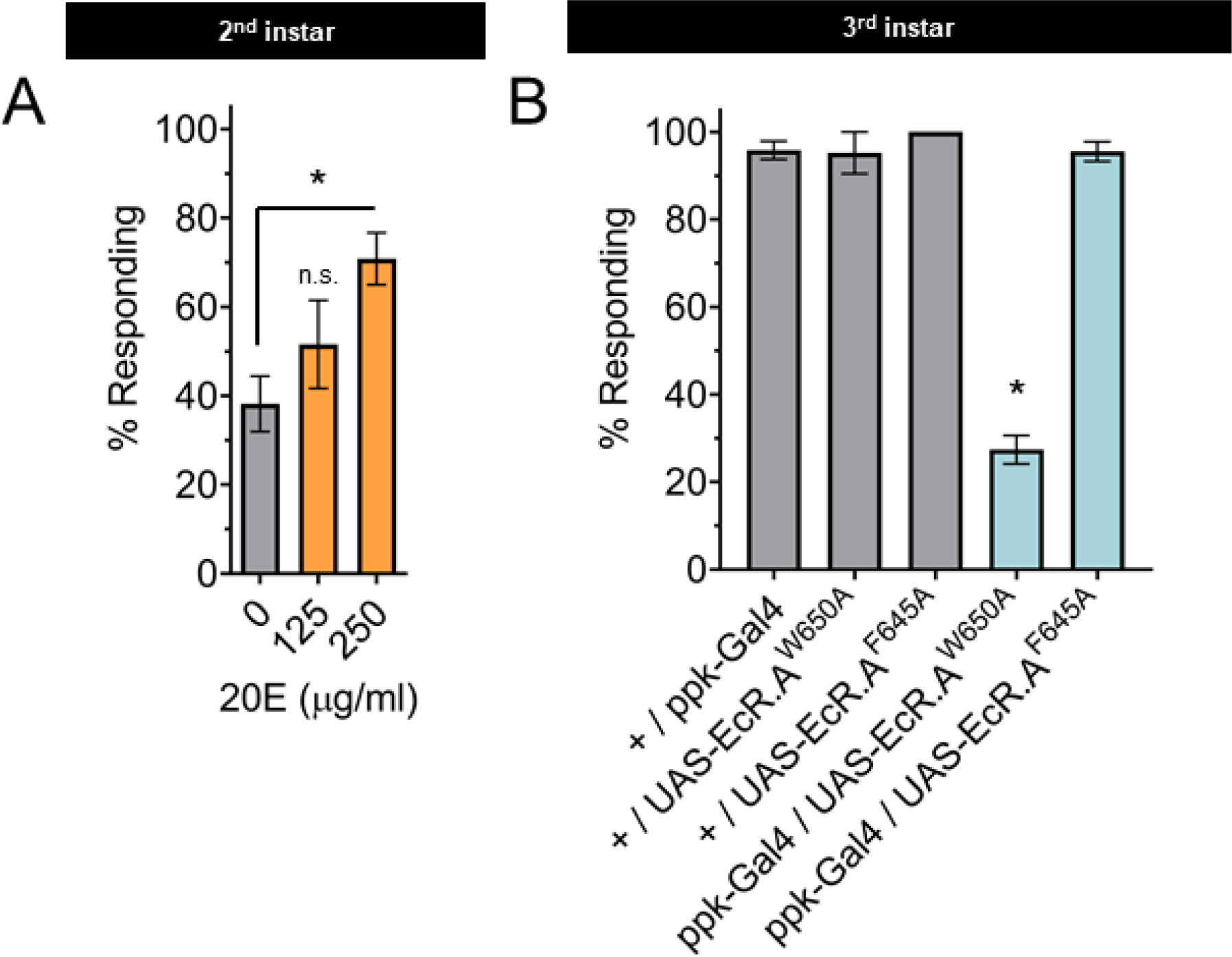
The nociceptive transition is ecdysone ligand activity dependent. (A) Percent of 2nd instar larvae population displaying nociceptive behavior when fed 20-hydroxyecdysone (20E). (B) Percent of population displaying nociceptive behavior when overexpressing ligand binding mutant EcR-A (ppk-Gal4/UAS-EcR-A -W650A), or co-activator mutant EcR-A (ppk-Gal4/UAS-EcR-A-F645A) with 42°C nociceptive probe. A: One-way Anova. * p < 0.05. n > 35 larvae for each treatment. B: One-way Anova. * p < 0.05. n = 2-4 staging replicates of 15-20 larvae were tested for each genotype.

The binding of ecdysone to EcR alters the recruitment of cofactors involved in derepression and activation of transcription. In the 3^rd^ instar, reduction of EcR-A expression reduces thermal nociception (McParland et al. 2015). Therefore, we tested the effects of disrupting the EcR-A domains responsible for either ligand binding or co-activator recruitment on nociception, by characterizing mutants with alanine substitution in either the ligand binding domain (EcR-A-W650A) or the co-factor recruitment domain (EcR-A-F645A) (Cherbas et al. 2003). Mutant forms of EcR-A can function in a dominant negative capacity by outcompeting the endogenous wildtype EcR-A thereby inactivating the endogenous receptor function (Cherbas et al. 2003). In this way, overexpression of these mutant EcR-A constructs can distinguish the requirements of co-activation (disrupted by F645A) or ligand induced derepression (disrupted by W650A). When assaying for a dominant negative mutant phenotype, we observed that expression of EcR-A-F645A in C4da neurons did not change nociceptive behavior in 3^rd^ instar larvae, while expression of EcR-A-W650A reduced the number of larvae exhibiting nociceptive behavior in 3^rd^ instar larvae (Figure 3B). Of the larvae that did exhibit nociceptive behavior, expression of EcR-A-W650A did not alter their response latency (Figure 3 – figure supplement 1A,C). We conclude that ecdysone ligand binding activity of EcR-A is required for nociception in 3^rd^ instar larvae.

TrpA1 knockout larvae lacking expression of all TrpA1 isoforms lose nociceptive behavior in response to thermal stimuli at 40-44°C, but are still able to respond to a 46°C probe with nociceptive behavior (Gu et al. 2019). Therefore, we sought to determine whether disruption of EcR function has effects specific to the probe temperature. Interestingly, EcR-A-W650A expression affected nociceptive behavior induced with the 42°C probe (Figure 3B) but not with the 46°C probe (Figure 3 – figure supplement 1B,C). It thus appears that EcR-A co-activator assembly is not required for 3^rd^ instar nociception, while ligand binding activity is required for 3^rd^ instar nociceptive behavior in response to a 42°C probe.

Having found that ecdysone activation of EcR-A is required for nociception, we sought to determine if overexpression of EcR-A in C4da neurons is sufficient to render 2^nd^ instar larvae precociously nociceptive. Since 2^nd^ instar larvae have a low thermal nociceptive response rate, we assayed for a gain of thermal nociceptive response in 2^nd^ instar larvae with EcR-A overexpression in their C4da neurons. Indeed, overexpression of EcR-A was sufficient to increase thermal nociceptive behavior in 2^nd^ instar larvae (Figure 4A,B). In contrast to the temperature specific effect of dominant negative mutant expression in the 3^rd^ instar, overexpression of EcR-A in the 2^nd^ instar was able to promote nociception at both 42°C and 46°C probe temperatures (Figure 4B, Figure – figure supplement 1B). Of the larvae which did display a nociceptive behavior, overexpression of EcR-A did not significantly change the latency of the nociceptive response as compared to 2^nd^ instar control larvae (Figure 4 – figure supplement 1A,C).

**Figure 4:**
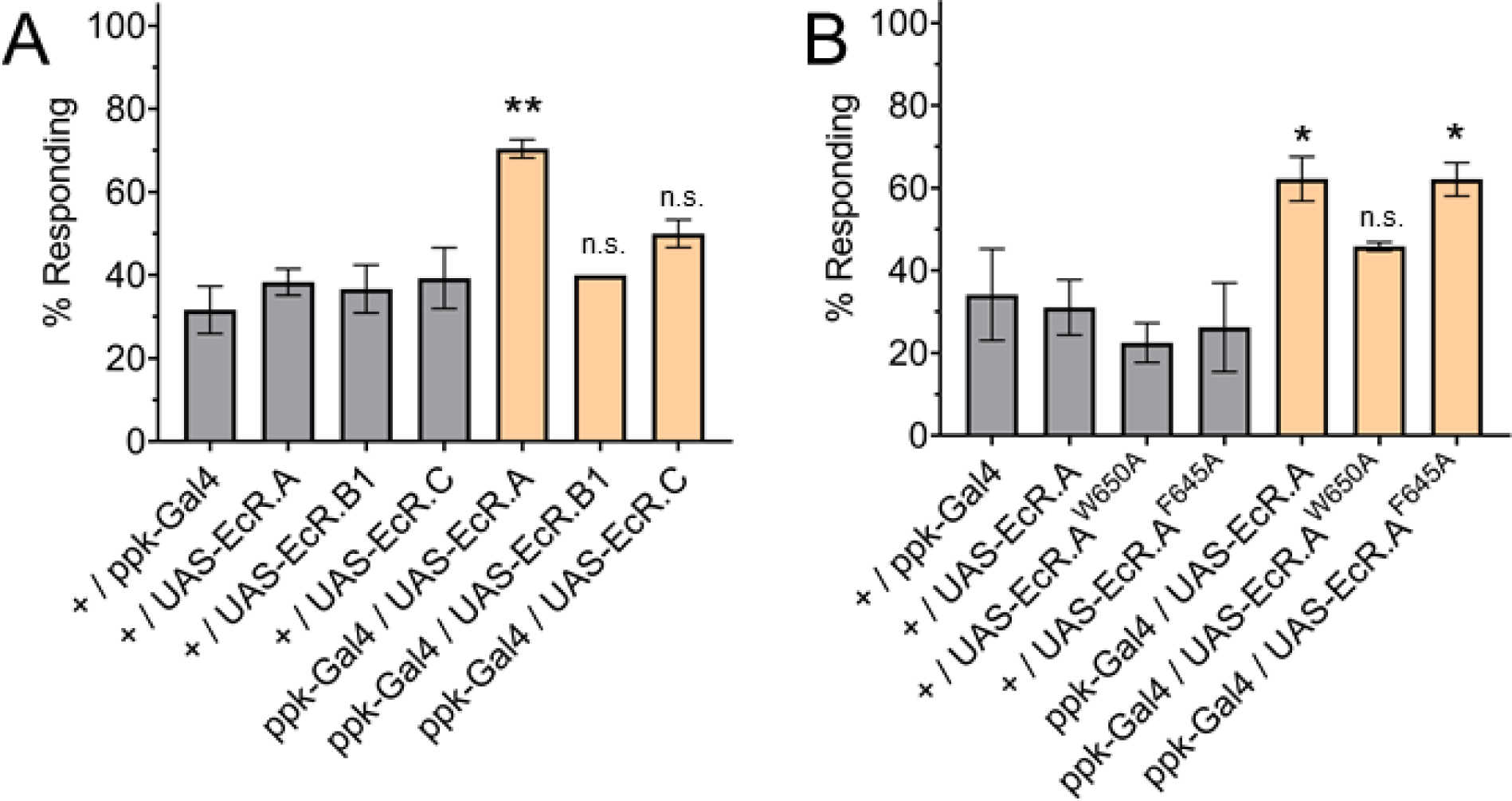
EcR-A overexpression in C4da neurons increases nociception in the 2^nd^ instar. **(A) Nociceptive** behavior in 2^nd^ instar larvae with EcR isoforms overexpressed in C4da neurons (B) Nociceptive behavior in 2^nd^ instar larvae with overexpression of EcR-A mutants. Ligand binding mutant EcR-A (ppk-Gal4/UAS-EcR-W650A), or co-activator mutant EcR-A (ppk-Gal4/UAS-EcR-F645A). 42°C nociceptive probe. One-way Anova with Tukey’s. * p < 0.05 ** p < 0.01. n = 2-4 staging replicates of 15-20 larvae were tested for each genotype.

Previous studies have found that overexpression of each EcR isoform can suppress expression of the other EcR isoforms (Schubiger et al. 2003). Additionally, overexpression of each EcR isoform could rescue loss of EcR function mutations in specific tissues (Cherbas et al. 2003). The difference between EcR isoforms is within the AF1 transcriptional activation domain (Figure 2 – figure supplement 1A). Therefore, we tested whether nociception in 2^nd^ instar larvae could be increased by overexpression in C4da neurons of EcR-A, EcR-B1 or EcR-C that contains only the sequences common to all isoforms and no AF1 transcriptional activation domain. We found that neither overexpression of EcR-B1 nor overexpression of EcR-C could increase nociception of 2^nd^ instar larvae (Figure 4A). Additionally, overexpression of these EcR isoforms did not alter nociception in the 3^rd^ instar (Figure 4 – figure supplement 2A,B). Given that expression of neither the domains common to all EcR isoforms, which are present in EcR-C, nor the AF1 domain of EcR-B1 could increase nociception in the 2^nd^ instar, we conclude that the AF1 transcriptional activation domain unique to EcR-A is required to increase nociception of 2^nd^ instar larvae.

Expression of *EcR-A* RNAi or EcR-B1-dominant negative mutant in the 3^rd^ instar larvae reduces the number of dendrite tips (Ou et al. 2008; McParland et al. 2015). Insensitivity to nociception is often associated with a reduction in dendrite structure, while hypersensitivity to nociception can be associated with both increased or decreased dendrite coverage (Honjo et al. 2016). We examined whether the EcR-A overexpression induced nociception was associated with changes of the dendrite branch number or arbor size. We found that EcR-A overexpression in the 2^nd^ instar reduced the number of dendrite tips (a measure of branch number) without significantly changing the area of the dendrite arbor (Figure 4 – figure supplement 2D-G). Our observation of decreased dendrite tip number with an associated increase in nociception is consistent with previously observed trends of decreased dendrite coverage in other nociceptive hypersensitive phenotypes (Honjo et al. 2016), further adding to the evidence that dendrite structure alone cannot predict nociceptive sensitivity.

Having found that ecdysone binding to EcR-A is required for 3^rd^ instar nociception, we sought to determine whether ligand binding is required for the precociously induced 2^nd^ instar nociception. We found that with overexpression of EcR-A-F645A, larvae were still precociously nociceptive as 2^nd^ instar larvae and they responded at a similar level as those overexpressing wildtype EcR-A (Figure 4B), suggesting that coactivator recruitment is not required for EcR-A induction of thermal nociception. In contrast, overexpression of the ligand binding mutant EcR-A-W650A did not facilitate precocious development of nociceptive behavior in 2^nd^ instar larvae to the level exhibited by larvae overexpressing wildtype EcR-A (Figure 4B). These effects of EcR mutant expression were observed at both 42°C and 46°C probe temperatures (Figure 4B, Figure – figure supplement 1B). Of those larvae which did display a nociceptive behavior, overexpression of EcR-A did not significantly change the latency of the nociceptive response (Figure 4 – figure supplement 1A,C). These findings reveal that thermal nociceptive development in the 2^nd^ instar larvae is sensitive to ecdysone regulation mediated by EcR-A via its ligand binding domain.

Together, these results demonstrate that EcR-A signaling is necessary and sufficient for the development of thermal nociceptor behavior. EcR-A acts in a cell autonomous manner in the C4da sensory neurons and requires the function of the ligand binding domain. Our results also suggest that co-activator recruitment is not required for thermal nociceptive development or maintenance. Additionally, our results suggest that in the 3^rd^ instar, the maintenance of ability to detect 46°C is independent of EcR-A ligand binding.

### Ecdysone receptor regulates *subdued* expression during larval thermal nociceptive development

Having found that the EcR-A AP1 domain and ligand binding are required for development of thermal nociception, we sought to determine whether any of the following nociceptive sensors are transcriptionally regulated by EcR-A. TrpA1 and Painless are transient receptor potential (TRP) channels required for thermal nociceptive behavior and nociceptive temperature-induced Ca^2+^ activity in C4da neurons (Tracey et al. 2003; Sokabe et al. 2008; Zhong et al. 2012; Terada et al. 2016; Gu et al. 2019). The TMEM16 family member Subdued is also required for thermal nociceptive behavior (Jang et al. 2015). To assess the scope of EcR-A regulation, we also included the degenerin/epithelial sodium channel (DEG/ENaC) channel Ppk1 that is required for ambient temperature driven behavior (Ainsley et al. 2008) and also functions along with Ppk26 in mechanical nociception (Gorczyca et al. 2014; Mauthner et al. 2014), as well as Piezo, which is involved in mechanical nociception but not thermal sensation in C4da neurons (Kim et al. 2012). In order to compare the changes in gene expression during the sensory switch in larval development, we conducted qRT-PCR with the nociceptive genes in GFP labeled C4da neurons isolated by Fluorescence Activated Cell Sorting (FACS) from control 2^nd^ and 3^rd^ instar larvae. We found decreased expression of *TrpA1*, *subdued*, *ppk1* and *ppk26* in 3^rd^ instar when compared to 2^nd^ instar C4da neurons. In contrast, *painless* expression increased while *piezo* expression did not significantly change from 2^nd^ to 3^rd^ instar C4da neurons (Figure 5A).

**Figure 5:**
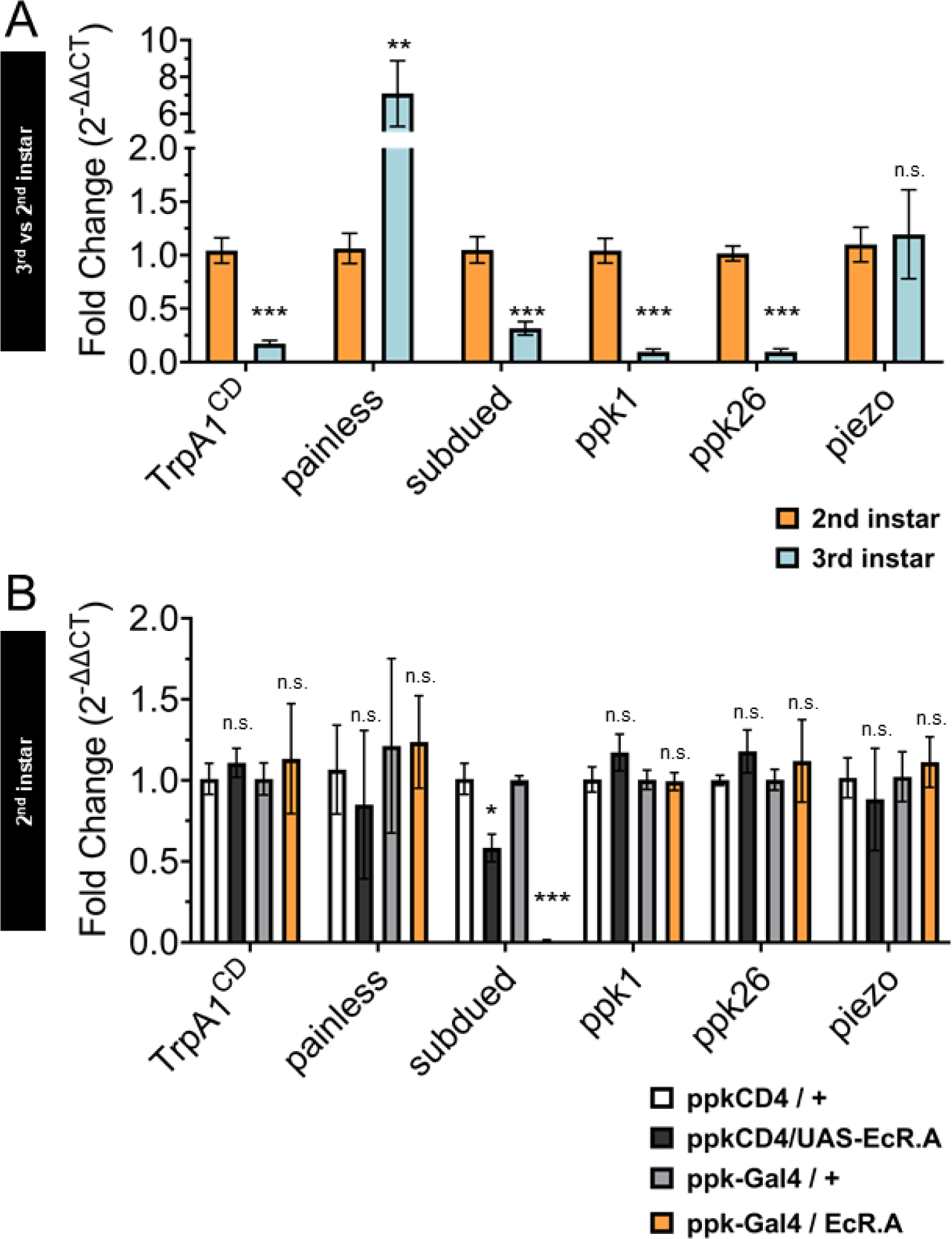
Ecdysone Receptor transcriptionally regulates *subdued* during the period of thermal nociceptive development. Expression of nociceptive genes measured by qRT-PCR from FACS purified C4da neurons. (A) C4da neurons isolated from 2^nd^ and 3^rd^ instar larvae. Expression in 3^rd^ instar neurons was normalized to 2^nd^ instar expression. (B) C4da neurons isolated from 2^nd^ instar larvae expressing wildtype EcR-A. Genotypic controls not expressing EcR-A (ppkCD4/UAS-EcR-A) were normalized to age matched neurons (ppkCD4/+). Expression in EcR-A expressing neurons (ppk-Gal4/UAS-EcR-A) was normalized to age matched control neurons without EcR-A expression (ppk-Gal4/+). Student’s t-test. * p < 0.05, *** p < 0.001.A: n = 8 and B: n = 3-4 FACS isolation/staging replicates for each age and genotype.

Next, we overexpressed EcR-A in C4da neurons and isolated these neurons with FACS from 2^nd^ instar larvae to measure expression of the nociceptive sensors. Comparing transcript levels of EcR-A-overexpressing C4da neurons to controls revealed that *subdued* expression was suppressed in EcR-A expressing C4da neurons as compared to that in control 2^nd^ instar C4da neurons (Figure 5B), thus mimicking the decrease during developmental transition from 2^nd^ to 3^rd^ instar. Notably, *subdued* stood out amongst the channel genes tested as the only one that displayed EcR-A regulation consistent with the developmental sensory switch.

While overexpression of EcR-A increased nociception in 2^nd^ instar larvae, overexpression of the ligand binding mutant EcR-A-W650A did not increase nociception in 2^nd^ instar but reduced nociception in the 3^rd^ instar (Figure 3 and 4). Having found that EcR-A overexpression specifically suppressed *subdued* expression, we sought to determine whether expression of the EcR-A dominant negative mutant has any effect on the level of *subdued* expression. When we overexpressed EcR-A-W650A in C4da neurons and then used FACS to isolate these neurons from either 2^nd^ instar or 3^rd^ instar larvae, we observed a reduction of *subdued* expression at both stages of larval development (Figure 5 – figure supplement 1A,B). Additionally, we found that expression EcR-A-W650A in 2^nd^ instar larvae significantly reduced *TrpA1*, *ppk26*, and *piezo* expression (Figure 5 – figure supplement 1A), while EcR-A-W650A expression in 3rd instar larvae significantly reduced *ppk1*, *ppk26*, and *piezo* expression (Figure 5 – figure supplement 1B).

Because *EcR-A* RNAi expression in C4da neurons has been shown to reduce nociception in the 3^rd^ instar, we also measured expression of the nociceptive sensors from FACS isolated 3^rd^ instar C4da neurons which expressed *EcR-A* RNAi. Expression of the *EcR-A* RNAi robustly reduced the presence of EcR-A protein in C4da neurons (Figure 5 – figure supplement 2A), but we found no significant difference in the expression of nociceptive sensor genes (Figure 5 – figure supplement 2B).

These data demonstrate that EcR-A regulates *subdued* expression. EcR-A overexpression can specifically suppress *subdued* expression in 2^nd^ instar larvae, as expected from the decrease of *subdued* expression in 3^rd^ instar larvae. We also found that inhibiting EcR-A ligand binding capacity with expression of EcR-A-W650A can reduce expression of more nociceptive genes than expression of wildtype EcR-A. EcR-A-W650A expression reduced *subdued* expression in the 2^nd^ instar, and further reduced expression of *subdued* in 3^rd^ instar larvae. We conclude that during the developmental period of the sensory switch, EcR-A regulates expression of *subdued*, and that ligand binding activity is necessary for maintaining the regulated expression of multiple nociceptor genes in the 3^rd^ instar.

### Reduction of *subdued* confers 2nd instar nociceptors with thermal and light sensitivity characteristic of 3^rd^ instar nociceptors

Our transcriptional data indicate that EcR-A overexpression specifically suppresses *subdued* expression in 2^nd^ instar C4da neurons. Therefore, we sought to determine whether *subdued* expression in 2^nd^ instar C4da neurons is responsible for the suppression of thermal nociceptive behavior in early larval development.

A knockout mutant of the *subdued* gene locus (*subdued^KO11^*) (Wong et al. 2013) eliminated *subdued* mRNA expression (Figure 6 – figure supplement 1A), but did not exhibit a significant change of nociceptive behavior in 2^nd^ instar larvae (Figure 6A). However, *subdued^KO11^* larvae were significantly smaller than age matched 2^nd^ instar control larvae (Figure 6 – figure supplement 1B), raising the possibility that pleiotropic effects of *subdued* loss of function mutation in the whole animal may alter the behavioral dynamics and mask the effect of mutation in the C4da neurons. Expression of a *subdued*-RNAi was effective at reducing *subdued* expression but did not eliminate all expression (Figure 6 – figure supplement 1C). Therefore, we tested the effect of expression of *subdued*-RNAi in C4da neurons, which did not reduce the size of 2^nd^ instar larvae (Figure 6 – figure supplement 1D). When *subdued*-RNAi was expressed specifically in the C4da neurons, 2^nd^ instar larvae had a greater nociceptive behavioral response than controls (Figure 6B). Thus, RNAi reduction of *subdued* expression in the C4da neurons caused a precocious thermal nociceptive response in 2^nd^ instar larvae, phenocopying EcR-A overexpression.

**Figure 6:**
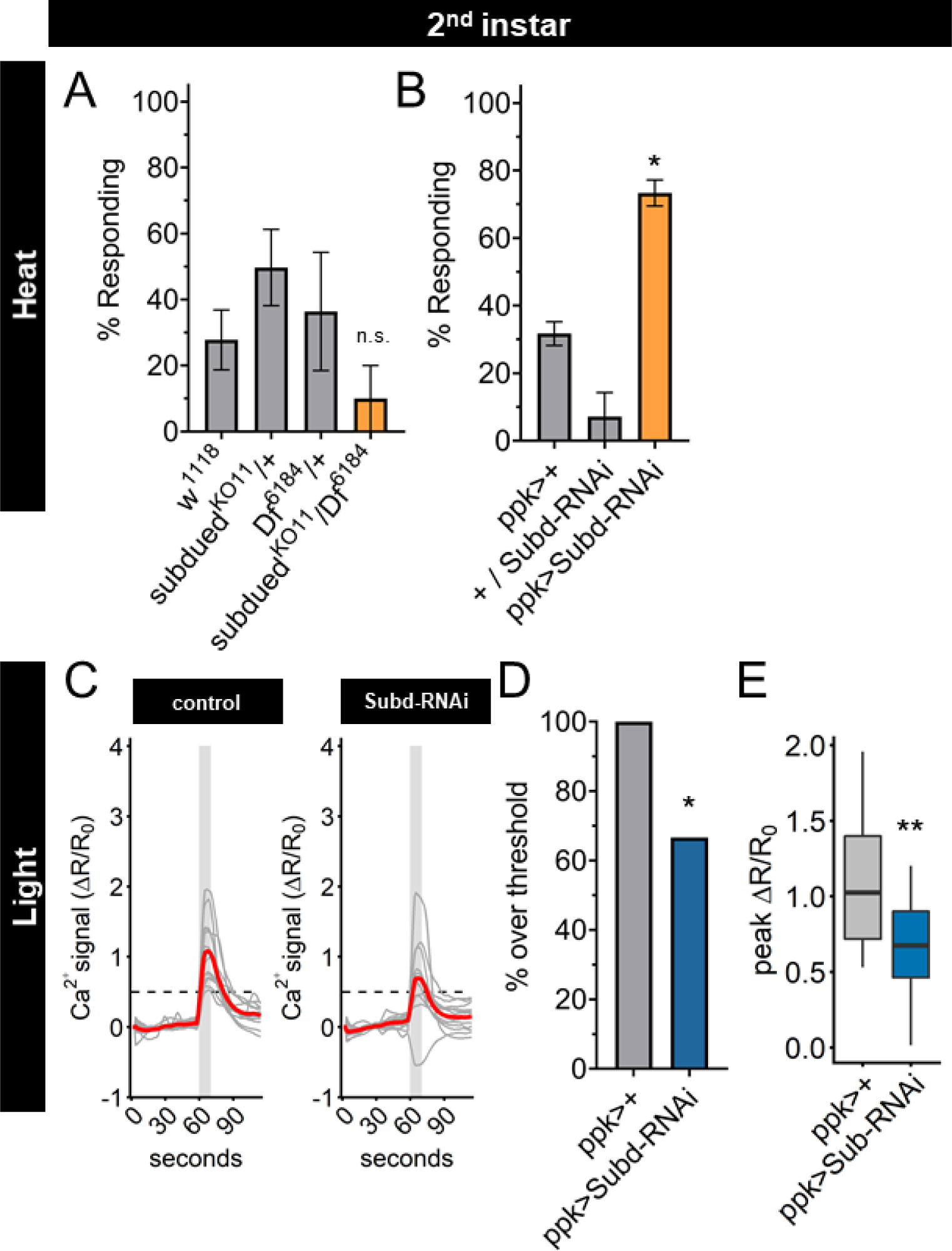
Subdued RNAi changes 2nd instar C4da neuron sensory response pattern to 3rd instar sensory response pattern. (A) 2^nd^ instar larvae with 42°C nociceptive probe. Percent of population displaying nociceptive behavior when mutant for Subdued. (B) 2^nd^ instar larvae with 42°C nociceptive probe. Percent of population displaying nociceptive behavior when expressing Subdued-RNAi (ppk-Gal4>Subd-RNAi v37472). (C) Individual traces (grey lines) and means (red lines) of Ca^2+^ activity calculated by ratiometric change from baseline. Grey column indicates period of stimulus. Dotted line indicates threshold level for % over threshold calculations. (D) Percent of soma with Ca^2+^ activity over the threshold level. (E) Peak Ca^2+^ activity during the periods of stimulus programs. A,B: One-way Anova. D: Fisher’s exact test, one-sided. E: Student’s t-test. A,B: n = 3-4 staging replicates of 15-20 larvae. C-E: ppk>+ n = 14 neurons, ppk>Subd-RNAi n = 12 neurons. * p < 0.05, ** p < 0.01.

Having found that UV light induced Ca^2+^ activity in C4da neurons decreased during larval development, we sought to determine whether Subdued is involved in the reduction of UV-A Ca^2+^ activity from 2^nd^ to 3^rd^ instar. We observed that expression of *subdued*-RNAi decreased both the number of actively responding neurons and the peak Ca^2+^ activity in C4da neurons during light stimulation of 2^nd^ instar larvae (Figure 6C-E).

Together these data demonstrate that reducing expression of *subdued* in 2^nd^ instar larvae is sufficient not only to increase thermal nociception but also to decrease the UV-A light response, thus enabling 2^nd^ instar C4da neurons to respond to thermal as well as light stimuli in the same manner as 3^rd^ instar C4da neurons. These effects of reduced *subdued* expression match the developmental progression of the sensory switch. We conclude that EcR-A regulation of the C4da sensory switch is mediated in part by reduction of *subdued* expression levels: *subdued* expression, owing to the EcR-A unliganded activity in 2^nd^ instar C4da neurons, is associated with a lack of thermal nociception, whereas suppression of *subdued* expression via EcR-A overexpression or *subdued*-RNAi in 2^nd^ instar C4da neurons causes precocious thermal nociception while reducing responsiveness to UV light (Figure 6 – figure supplement 1E).

## Discussion

The impact of peripheral nervous system (PNS) development on behavior is underscored by the recent finding that defects in the PNS are sufficient to produce symptoms in mouse models of autism spectrum disorders (Orefice et al. 2016). The role of hormonal regulation of sensory systems during periods of postnatal development has also been highlighted by studies of growth hormone regulation of thermal sensitivity (Liu et al. 2017; Ford et al. 2019). Considering that sensory neurons progress through transcriptionally distinct identities during development (Sharma et al. 2020), characterizing how the specific changes in receptors or channels that produce these developmentally specific outcomes is of prominent importance for understanding the development of neurological disease.

In this study we address how a nociceptive behavior is temporally acquired during development. We demonstrate how steroid hormone regulation of nociceptor activity in a class of sensory neurons in the PNS adjusts the developing larval response to nociceptive stimuli. Heat induced nociceptive behavior in *Drosophila melanogaster* larvae arises during the final (3^rd^) instar of development (Sulkowski et al. 2011). Here, we show that this behavioral transition reflects a nociceptive sensory switch through regulation of Subdued by the steroid hormone ecdysone and the ecdysone receptor isoform EcR-A.

While previous studies have focused on EcR localization in the 3^rd^ instar and pupal stages of C4da neurons, we show EcR is present in both the nucleus and cytoplasm in the 2^nd^ instar, followed with an increase of EcR nuclear localization 8 hrs into the 3^rd^ instar (Figure 2). Previous work has shown that loss of EcR-A reduces thermal nociception in the 3^rd^ instar (McParland et al. 2015). While we did not find detectable transcriptional changes in nociceptor genes in 3^rd^ instar C4da neurons during *EcR-A* RNAi expression, we found broad suppression of nociceptor genes during EcR ligand binding mutant (W650A) expression (Figure 5 – figure supplement 2). Through behavioral and transcriptional analysis, we show that EcR-A nociception regulation is ligand dependent, by demonstrating that increased ecdysone titer and overexpression of ligand-competent EcR both accelerate nociception in the 2^nd^ instar (Figure 3 and Figure 4). Ligand binding of EcR leads to widespread epigenetic changes through release of co-repressors and recruitment of co-activators (Uyehara et al. 2017; Uyehara and McKay 2019). The precocious activation of nociception in the 2^nd^ instar requires the unique AF1 domain of the isoform EcR-A, as other isoforms lacking this domain do not increase nociception (Figure 4A). The AF1 domain of EcR-A is less activating and more repressive then the EcR-B1 AF1 domain (Dela Cruz et al. 2000; Mouillet et al. 2001; Hu et al. 2003). It thus appears that ligand dependent derepression could be the action which promotes nociception. Moreover, expression of the EcR co-activator mutant (F645A) phenocopies expression of wildtype EcR in its ability to promote nociception (Figure 4B), suggesting that in contrast to co-activator recruitment, preservation of the ability to remove co-repressors is required for nociception. This regulation of repression may account for the ability of the EcR ligand binding mutant (W650A) to suppress multiple nociceptive genes including *subdued* in the 2^nd^ and 3^rd^ instar. The release of co-repressors could allow other transcription factors to gain access to response elements which suppress *subdued*, or directly promote transcription of a repressor of *subdued*. How derepression leads to the suppression of *subdued* is an intriguing open question.

For 3^rd^ instar larval nociception, our data suggest there are both ligand dependent and ligand independent EcR-A pathways. We find 3^rd^ instar expression of EcR-A ligand binding mutant to have a temperature specific phenotype. Expression of the EcR-A ligand binding mutant inhibits nociception at 42°C, but does not reduce nociception at 46°C (Figure 3B, Figure 3 – figure supplement 1B). This temperature specific effect is reminiscent of the phenotype of TrpA1 mutant larvae, which lose nociceptive behavior at probe temperatures of 44°C and lower, but are still responsive to 46°C stimuli (Gu et al. 2019). We found that expression of EcR-A ligand binding mutant reduced *subdued* expression in the 3^rd^ instar even more than during the developmental transition (Figure 5A,C), suggesting that this loss of *subdued* expression may contribute to the phenotype at 42°C but not at 46°C. Consistent with this possibility, we observed that Subdued-RNAi was able to increase nociception in 2^nd^ instar larvae while Subdued knockout mutation did not increase nociception. Together these results suggest that the amount of *subdued* expression is important for promoting or inhibiting nociception. Furthermore, there are Subdued and EcR-A ligand-independent pathways which promote nociception in the 3^rd^ instar. We observe alteration of *TrpA1*, *painless*, *ppk1 and ppk26* expression during development but not as a result of EcR overexpression (Figure 5). These separate pathways may involve mechanisms of regulating temperature specificity in C4da neurons.

Subdued has homology to the Calcium activated Chloride Channels (CaCC) of the mammalian TMEM16 family, with channel activity similar to both TMEM16A and TMEM16F (Wong et al. 2013; Le et al. 2019). The effect of CaCC on neural activity is dependent on the intracellular Cl^-^ concentration ([Cl^-^]i) (Berg et al. 2012). When [Cl^-^]i is high, elevation of internal Ca^2+^ level activates CaCCs and causes Cl^-^ efflux, leading to excitation. In contrast, when [Cl^-^]i is low, Ca^2+^ activation of CaCCs causes an influx of Cl^-^ which has an inhibitory effect on neuronal excitability. Differential expression of symporters and co-transporters in the CNS and PNS, as well as in immature and mature neurons, is important for creating the inhibitory or excitatory effects of CaCCs. In 3^rd^ instar larvae, Cl^-^ efflux has been observed during thermal nociceptive stimulation of C4da neurons (Onodera et al. 2017). However, Onodera and colleagues found that loss of Subdued did not significantly change the magnitude of the Cl^-^ efflux during heat stimulus, suggesting other channels are responsible for the Cl^-^ efflux in 3^rd^ instar C4da neurons. The direction of Cl^-^ movement in the 2^nd^ instar remains to be determined, and it is conceivable that Subdued could produce an inhibitory influx of Cl^-^ in the 2^nd^ instar. Characterizing Cl^-^ homeostasis during C4da neuron development and its impact on nociception will be of interest in future studies.

Subdued has previously been found to regulate thermal nociception in 3^rd^ instar larvae (Jang et al. 2015). Jang and colleagues found the strongest decrease of nociception in the 3^rd^ instar when expression of *subdued* was reduced in all dendritic arborization neurons. Additionally, when *subdued* was overexpressed in C4da neurons the number of 3^rd^ instar larvae that responded to nociceptive heat did not significantly change. Interesting, when *subdued* was overexpressed with a subdued-Gal4, the number of fast responding 3^rd^ instar larvae increased (Jang et al. 2015). This suggests there may be a substantial thermal nociception role for Subdued in dendritic arborization neurons other than C4da neurons during 3^rd^ instar. Our data suggest that as C4da neurons develop from 2^nd^ to 3^rd^ instar, the ability of Subdued to influence detection of stimuli may change. We find that thermal nociception is enhanced by reduction of *subdued* expression in 2^nd^ instar C4da neurons, via either EcR-A overexpression or Subdued-RNAi knockdown. Additionally, we find that expression of the EcR-A-ligand mutant reduces *subdued* expression even further in 3^rd^ instar C4da neurons, and that this is associated with a loss of nociception. The greater amount of *subdued* expression in the 2^nd^ instar may mean that Subdued could have a greater impact on Cl^-^ regulation during the 2^nd^ instar. This mechanism of Subdued activity would be dependent on the intracellular Cl^-^ concentration created by symporters and co-transporters. Therefore, identifying regulators of Cl^-^ concentration during the development of C4da neurons and their impact on nociception will be of interest in future studies.

Our work adds to the evidence that the developmental transition of thermal nociception represents a shift in sensory state rather than a change in sensory system construction: 1) optogenetics can stimulate aversive behavior in both 2^nd^ and 3^rd^ instar C4da neurons (Sulkowski et al. 2011), 2) TrpA1 misexpression can confer nociceptive behavior in both 2^nd^ and 3^rd^ instar larvae exposed to innocuous temperatures (Luo et al. 2017), 3) UV-A stimulation induces greater Ca^2+^ response in 2^nd^ instar than in 3^rd^ instar C4da neurons (Figure 1), and 4) reduction of *subdued* expression in 2^nd^ instar renders C4da neurons responsive to thermal nociceptive stimuli (Figure 6).

Recent findings have highlighted the occurrence of developmentally timed modulation of sensory state in both larvae and adult *Drosophila*. A sensory switch in thermotaxis behavior of *Drosophila* larvae is regulated by transcriptional regulation of thermoreceptors (Tyrrell et al. 2021). In adults, male courtship behavior is modulated through olfactory neuron pheromone sensitivity and hormone-mediate chromatin reprograming (Zhao et al. 2020). Our study highlights that temporally programed sensory switches may be a widely used developmental mechanism to match behavioral outcomes with life history events.

We speculate that the nociceptive shift in sensory state is an important mechanism for matching sensory system function with life history transitions. *Drosophila melanogaster* larvae do not develop robust thermal nociception until the second half of the larval phase. This means that for the first half of larval life, larvae do not have thermal nociception behavior. In contrast, nociceptor activity to UV-A light decreases from the 2^nd^ to 3^rd^ instar, suggesting that nociceptive sensitivity to short wavelength light decreases during the second half of the larval phase. This transition correlates with larval behavioral changes that emerge during the 3^rd^ instar, such as cessation of feeding beneath the surface and wandering above ground (Wegman et al. 2010). Larvae are transitioning from predominantly feeding behavior in the 2^nd^ instar, often shielded from light and high temperatures, to movement away from food in search of pupation sites in the 3^rd^ instar. In a natural habitat, this transition can mean leaving the food source, which necessitates greater exposure to heat, increasing the risk of predation and desiccation (Ballman et al. 2017). This sensory switch may function as a developmental strategy, shaping behavioral outcomes as animals encounter changing environments, thus increasing survival.

## Materials and Methods

### Drosophila melanogaster stocks and culturing

Detailed stock genotypes and sources are listed in Table 1. X chromosome genotypes are simplified: male and female larvae are pooled together in test populations, and the origin of miniwhite alleles could be mixed. Control genotypes are crossed to w^1118^ (v60000) unless otherwise noted.

**Table 1:** *Drosophila melanogaster* stocks used in this study

| Figure | Genotype | Source |
| --- | --- | --- |
| 1A and S1-1 | ppk-tdGFP<sup>1b</sup>,UAS-Dcr2/+;ppk-Gal4<sup>1a</sup>/+ | (Han et al. 2011) |
| 1C-F | ppk-Gal4<sup>vk37</sup>/+;UAS-GCaMP6(s),UAS-tdTom/+ | BDSC 42749, (Han et al. 2011) |
| 2A-C, S2-1B, 3A | ppk-tdGFP<sup>1b</sup>,UAS-Dcr2/+;ppk-Gal4<sup>1a</sup>/+ |  |
| 3B and S3-1A-C | ppk-tdGFP<sup>1b</sup>,UAS-Dcr2/+;ppk-Gal4<sup>1a</sup>/+ |  |
|  | P{UAS-EcR.A.W650A}TP5/+ | BDSC 9451 |
|  | P{UAS-EcR.A.F645A}TP2/+ | BDSC 9452 |
|  | ppk-tdGFP<sup>1b</sup>,UAS-Dcr2/P{UAS-EcR.A.F645A}TP2;ppk-Gal4<sup>1a</sup>/+ |  |
|  | ppk-tdGFP<sup>1b</sup>,UAS-Dcr2/P{UAS-EcR.A.W650A}TP5;ppk-Gal4<sup>1a</sup>/+ |  |
| 4A and S4-2A,B | ppk-Gal4<sup>vk37</sup> | (Han et al. 2011) |
|  | P{UAS-EcR.A}3a/+ | BDSC 6470 |
|  | P{UAS-EcR.B1}3b/+ | BDSC 6469 |
|  | P{UAS-EcR.C}TP1-4/+ | BDSC 6868 |
|  | ppk-Gal4<sup>vk37</sup>;P{UAS-EcR.A}3a |  |
|  | ppk-Gal4<sup>vk37</sup>; P{UAS-EcR.B1}3b |  |
|  | ppk-Gal4<sup>vk37</sup>;P{UAS-EcR.C}TP1-4 |  |
| 4B and S4-1A-G | ppk-tdGFP<sup>1b</sup>,UAS-Dcr2/+;ppk-Gal4<sup>1a</sup>/+ |  |
|  | P{UAS-EcR.A}3a/+ |  |
|  | P{UAS-EcR.A.F645A}TP2/+ |  |
|  | P{UAS-EcR.A.W650A}TP5/+ |  |
|  | ppk-tdGFP<sup>1b</sup>,UAS-Dcr2/+;ppk-Gal4<sup>1a</sup>/P{UAS-EcR.A}3a |  |
|  | ppk-tdGFP<sup>1b</sup>,UAS-Dcr2/P{UAS-EcR.A.F645A}TP2;ppk-Gal4<sup>1a</sup>/+ |  |
|  | ppk-tdGFP<sup>1b</sup>,UAS-Dcr2/P{UAS-EcR.A.W650A}TP5;ppk-Gal4<sup>1a</sup>/+ |  |
| 5A | ppk-tdGFP<sup>1b</sup>,UAS-Dcr2/+;ppk-Gal4<sup>1a</sup>/+ |  |
| 5B | ppk-CD4- tdGFP<sup>1b</sup> /+ |  |
|  | ppk-CD4- tdGFP<sup>1b</sup> /+;P{UAS-EcR.A}3a/+ |  |
|  | ppk-tdGFP<sup>1b</sup>,UAS-Dcr2/+;ppk-Gal4<sup>1a</sup>/+ |  |
|  | ppk-tdGFP<sup>1b</sup>,UAS-Dcr2/+;ppk-Gal4<sup>1a</sup>/P{UAS-EcR.A}3a |  |
| S5-1A,B | ppk-CD4- tdGFP<sup>1b</sup> /+ |  |
|  | ppk-CD4- tdGFP<sup>1b</sup> / P{UAS-EcR.A.W650A}TP5 |  |
|  | ppk-tdGFP<sup>1b</sup>,UAS-Dcr2/+;ppk-Gal4<sup>1a</sup>/+ |  |
|  | ppk-tdGFP<sup>1b</sup>,UAS-Dcr2/P{UAS-EcR.A.W650A}TP5;ppk-Gal4<sup>1a</sup>/+ |  |
| S5-2A,B | ppk-tdGFP<sup>1b</sup>,UAS-Dcr2/+;ppk-Gal4<sup>1a</sup>/+ |  |
|  | ppk-CD4- tdGFP<sup>1b</sup> /+;P{w[+mC]=UAS-EcR.A.dsRNA}91/+ | BDSC 9328 |
|  | ppk-tdGFP<sup>1b</sup>,UAS-Dcr2/+;ppk-Gal4<sup>1a</sup>/P{w[+mC]=UAS-EcR.A.dsRNA}91 |  |
| 6A, S6-1A-B | Subdued<sup>KO11</sup>/+ | (Wong et al. 2013) |
|  | Df(3R)Exel6184, P{w[+mC]=XP-U}Exel6184 | BDSC 7663 |
|  | Subdued<sup>KO11</sup>/Df(3R)Exel6184, P{w[+mC]=XP-U}Exel6184 |  |
| 6B, S6-1D | ppk-tdGFP<sup>1b</sup>,UAS-Dcr2/+;ppk-Gal4<sup>1a</sup>/+ |  |
|  | P{GD3674}v37472/+ | VDRC 37472 |
|  | ppk-tdGFP<sup>1b</sup>,UAS-Dcr2/+;ppk-Gal4<sup>1a</sup>/P{GD3674}v37472 |  |
| S6-1C | P{GD3674}v37472/+ | VDRC 37472 |
|  | Actin-Gal4/+; Subdued<sup>KO2</sup>/+ | (Wong et al. 2013) |
|  | Actin-Gal4/+; Subdued<sup>KO2</sup>/ P{GD3674}v37472 |  |
| 6-1C-E | ppk-Gal4<sup>vk37</sup>/+;UAS-GCaMP6(s),UAS-tdTom/+ | BDSC 42749, (Han et al. 2011) |
|  | ppk-Gal4<sup>vk37</sup>/+;UAS-GCaMP6(s),UAS-tdTom/P{GD3674}v37472 |  |

All experimental crosses were raised at 25°C and 60% humidity, with a 12hr light-dark cycle, and fed cornmeal-molasses media. Larval development was synchronized by staged egg collection on grape agar, followed by synchronized collection of newly hatched L1 larvae. 2^nd^ instar larvae were assayed between 26-32 hrs after L1 larval hatching and 3^rd^ instar larvae were collected as pre-pupariation wandering larvae. All assays were performed during the dark light cycle.

### Nociceptive Behavior Assay

Nociceptive stimulation was performed as previously described (Babcock et al. 2009). Larvae were briefly rinse in dH20 and allowed to acclimatize for 1 minute on a vinyl sheet. The larvae were kept moist and a temperature-controlled heat probe (ProDev Engineering, TX) was used to apply heat between the larval body segments A3-A6 of forward crawling larvae. Nociceptive behavior was defined as at least one complete 360° roll. The time to complete one roll (latency) was measured up to a 20 second cutoff. Each larva was presented with a single stimulus to avoid habituation. Latency was calculated for each biological replicate as the time at which greater than or equal to 50% of the responding population (non-responding larvae we excluded) had responded to the stimulus.

### 20-hydroxyecdysone feeding

For food supplemented with 20-hydroxyecdysone (20E) (Sigma, H5142), 20E was dissolved in 100% ethanol and mixed with room temperature cornmeal-molasses media. An equivalent volume of ethanol alone was mixed with media as a control. Larvae were transferred to supplemented media 48hrs AEL and assayed for nociceptive behavior 8 hours later.

### Ultraviolet light response calcium imaging

C4da response to UV-A stimulation was measured as previously described (Yadav et al. 2019). A Leica SP5 confocal microscope with resonance scanner was used with the 20x oil immersion objective and 16x optical magnification. Imaging was done with 512 x 512 resolution and a slice thickness of 5 μm (20 Z slices) at 1.201 fps. A 405 laser line (50mW) was used at 100% laser power. Neurons were imaged for 60 seconds before a 10 second UV exposure. Z-stacks were acquired in the GFP, RFP, an UV channels. The ratiometric signal was quantified as described for thermal Ca^2+^ imaging.

### Cell purification and qRT-PCR

C4da neurons were isolated by dissecting 30-40 larvae in 1x PBS on ice. To increase C4da neuron concentration in the final cell suspension, the larval body wall was inverted and the CNS, imaginal discs, gut, and fatbody were removed. After dissection cell suspensions were prepared by vortexed with 1.5μl 1x Liberase TM (Roche, LIBTM-RO) in 500 μl cold PBS. To dissociate cells samples were incubated for 3 periods at 25°C at 1,000 RPM, triturating 10 times with a glass pipette between each incubation. Suspensions were strained through a 40μm cell strainer (Fisher Scientific) and brought to 1.5ml with Schneider’s media. 1μl ethidium homodimer-1 (ThermoFisher, L3224) was added before sorting to mark dead cells. C4da neurons were isolated by fluorescence-activated cell sorting (FACS) with an Aria II (Becton Dickinson). GFP+ nonautofluorescent RFP-events were sorted into 20μl lysis buffer (Thermofisher, KIT0204) and immediately frozen on dry ice. 100-1000 cells were isolated per dissection replicate. RNA was isolated with PicoPure RNA isolation kit (Thermofisher, KIT0204), cDNA was synthesized with Applied Biosystems High-Capacity cDNA Reverse Transcription Kit (ThermoFisher, 4368813), and qRT-PCR was performed using SYBR green (ThermoFisher A25742) with a QuantStudio7 (ThermoFisher). Relative expression was calculated using the detla-delta Ct method with the housekeeping gene eEF1α2 normalized to the mean Ct of the genotypic and isolation controls.

*Subdued* expression in knockout mutant and RNAi expressing larvae was measured using RNeasy Mini Kit (Qiagen 74104) isolation from 3-4 larvae per genotype. cDNA syntheses and qRT-PCR were performed as described FACS isolated cells.

### Primers

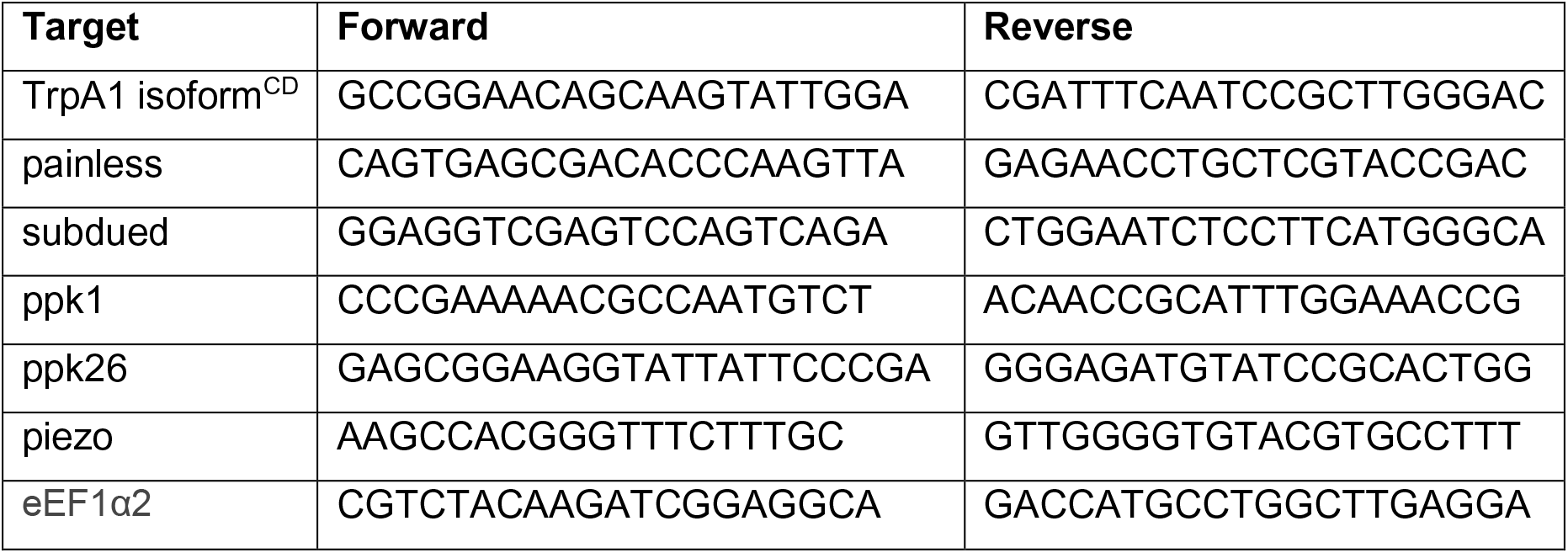

### Immunohistochemistry

Larval ages were staged by L1/L2 molt for 2^nd^ instar or L2/L3 molt for 3^rd^ instar. Larvae were filleted in PBS on ice and fixed for 10 min at room temperature in 4% paraformaldehyde. Samples were blocked with 5% goat serum for 2 hrs at room temperature and then incubated with either EcR-common DDA2.7, EcR-A 15G1a, or EcR-B1 AD4.4 (Developmental Studies Hybridoma Bank) in blocking solution (1:200) overnight at 4°C. After rinse, samples were incubated with secondary antibody Alexa Fluor 555 (Invitrogen A-2142A) in blocking solution (1:500) for 2 hrs at room temperature before 10 min staining with DAPI (1:10,000) and mounting in Vectashield (Vector Laboratories). Mean fluorescence was measured with Fiji (https://imagej.net/fiji) in traces of the nuclei and cytoplasm (traces of cytoplasm excluded C4da neuron nuclei and an neighboring nuclei) and adjusted for area before calculating the ratio of fluorescence in the nuclease and cytoplasm of C4da neurons.

### Larval size measurement

Larvae were collected in 75μl PBS and placed in an 80°C thermo-block for 10 min. The turgid larvae were then placed on a microscope slide, imaged, and larval body area was measured with Fiji (https://imagej.net/fiji).

### Statistical Analysis

Statistical tests were done with GraphPad Prism 8.3.0 (GraphPad Software) or R (version 3.5.2, R Core Team 2018).

## Acknowledgements

We thank: Tun Li and Susan Younger for experimental consultation, Caitlin O’Brien, Han-Hsuan Liu, Ke Li, Maja Petkovic, Beverly Piggott, and Rebecca Jaszczak for editorial advice. Stocks obtained from the Bloomington Drosophila Stock Center (BDSC, NIH P40OD018537) and Vienna Drosophila Resource Center (VDRC, www.vdrc.at) were used in this study. DDA27 was obtained from the from the Developmental Studies Hybridoma Bank (DSHB). Research reported in this publication was supported by the National Institutes of Health: National Institute of General Medical Sciences F32GM130019 (JSJ) and National Institute of Neurological Disorders and Stroke R35NS097227 (YNJ). YNJ and LYJ are investigators at the Howard Hughes Medical Institute.

---

**Figure 1 – figure supplement 1:**
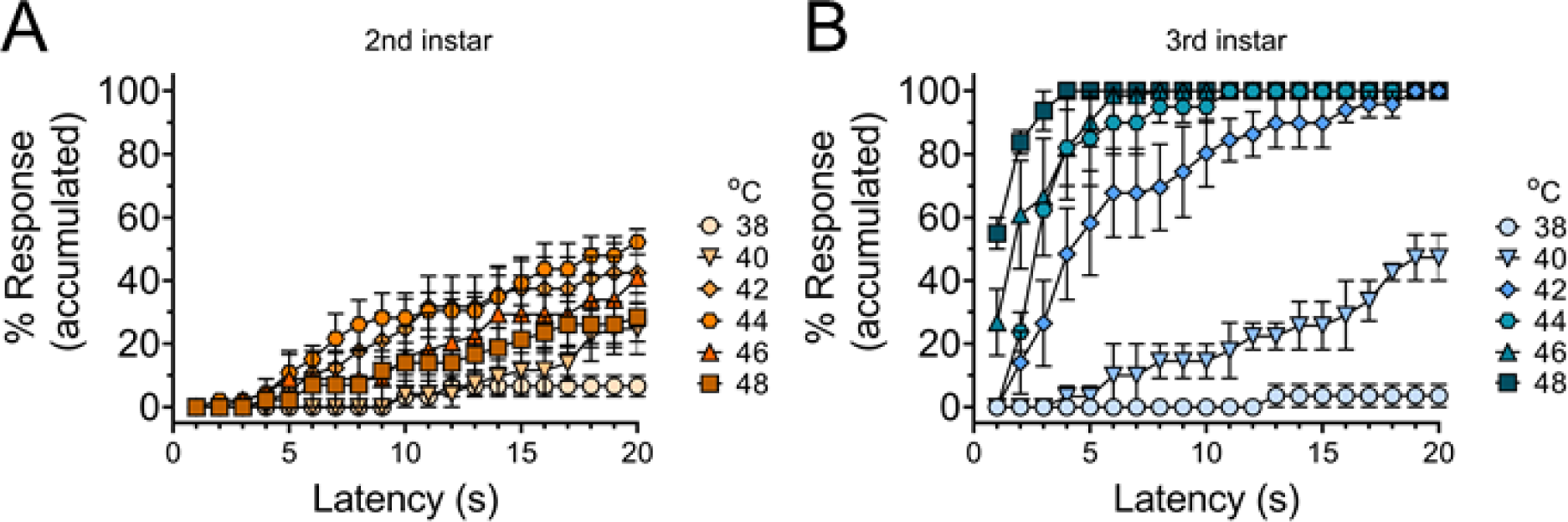
Thermal nociceptive sensitivity increases across temperatures from 2^nd^ to 3rd instar. Accumulated curve of percentage larvae that will have exhibited the nociceptive behavior by the length of time experiencing the stimulus (latency in seconds). (A) Percentage of 2^nd^ instar larvae that will have had the nociceptive behavior at the latency response time. (B) Percentage of 3^rd^ instar larvae that will have had the nociceptive behavior at the latency response time. Quantification of data from Figure 1A. n = 2-4 staging replicates of 15-20 animals were tested for each age and temperature.

**Figure 2 – figure supplement 1:**
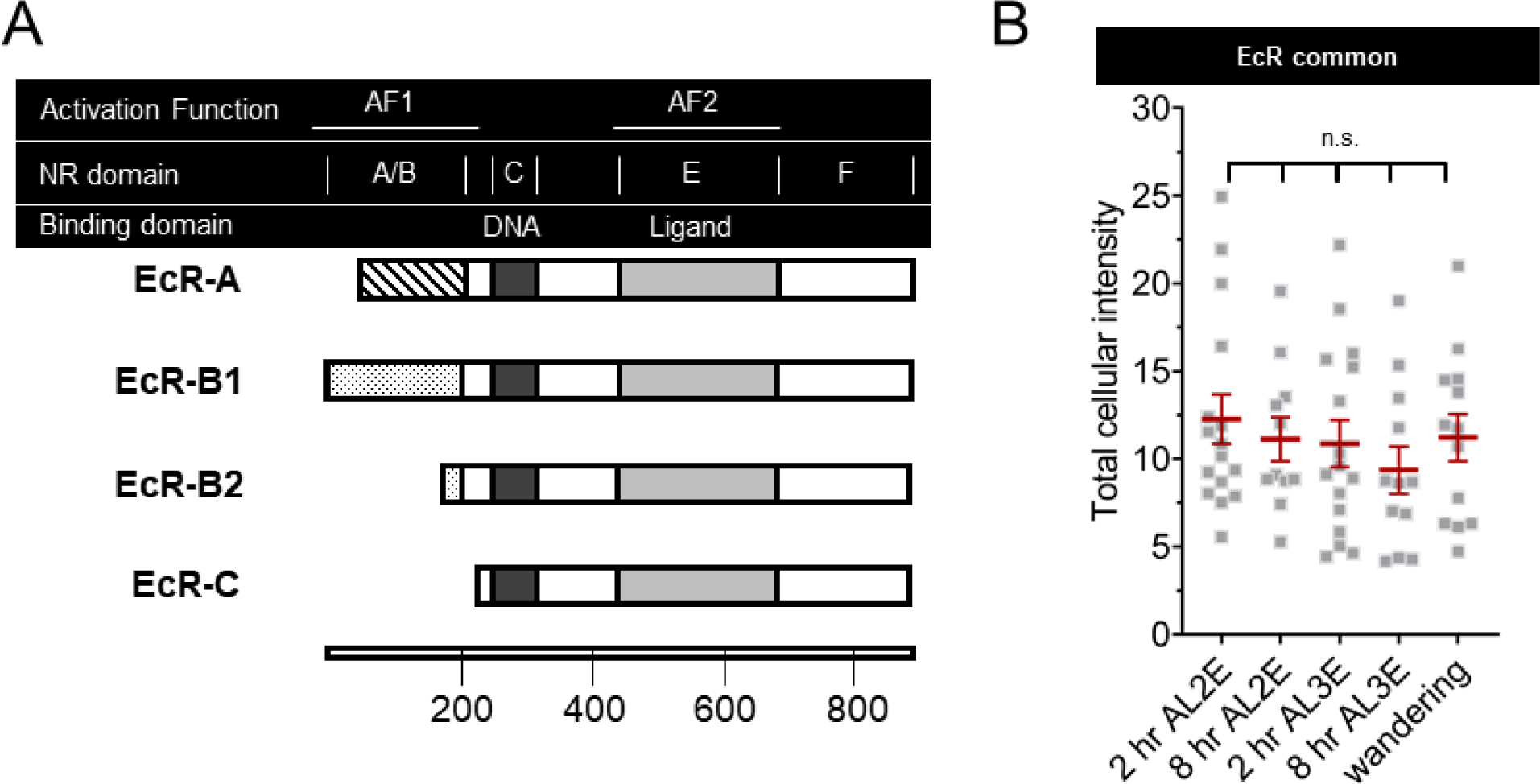
Total Ecdysone receptor remains constant in C4da neuron early during the 2^nd^ and 3^rd^ instar. (A) Ecdysone Receptor domain structures. Activation Function (AF) and Nuclear Receptor (NR) domains along with DNA (dark bar) and Ligand (light bar) binding domains. All EcR isoforms have identical sequences except for the A/B NR domain which contains AF1 function. Scale in amino acid residues. (B) EcR immunohistochemistry with EcR-common antibody quantifying intensity in both the nucleus and cytoplasm. One-way Anova with Bonferroni.

**Figure 3 – figure supplement 1:**
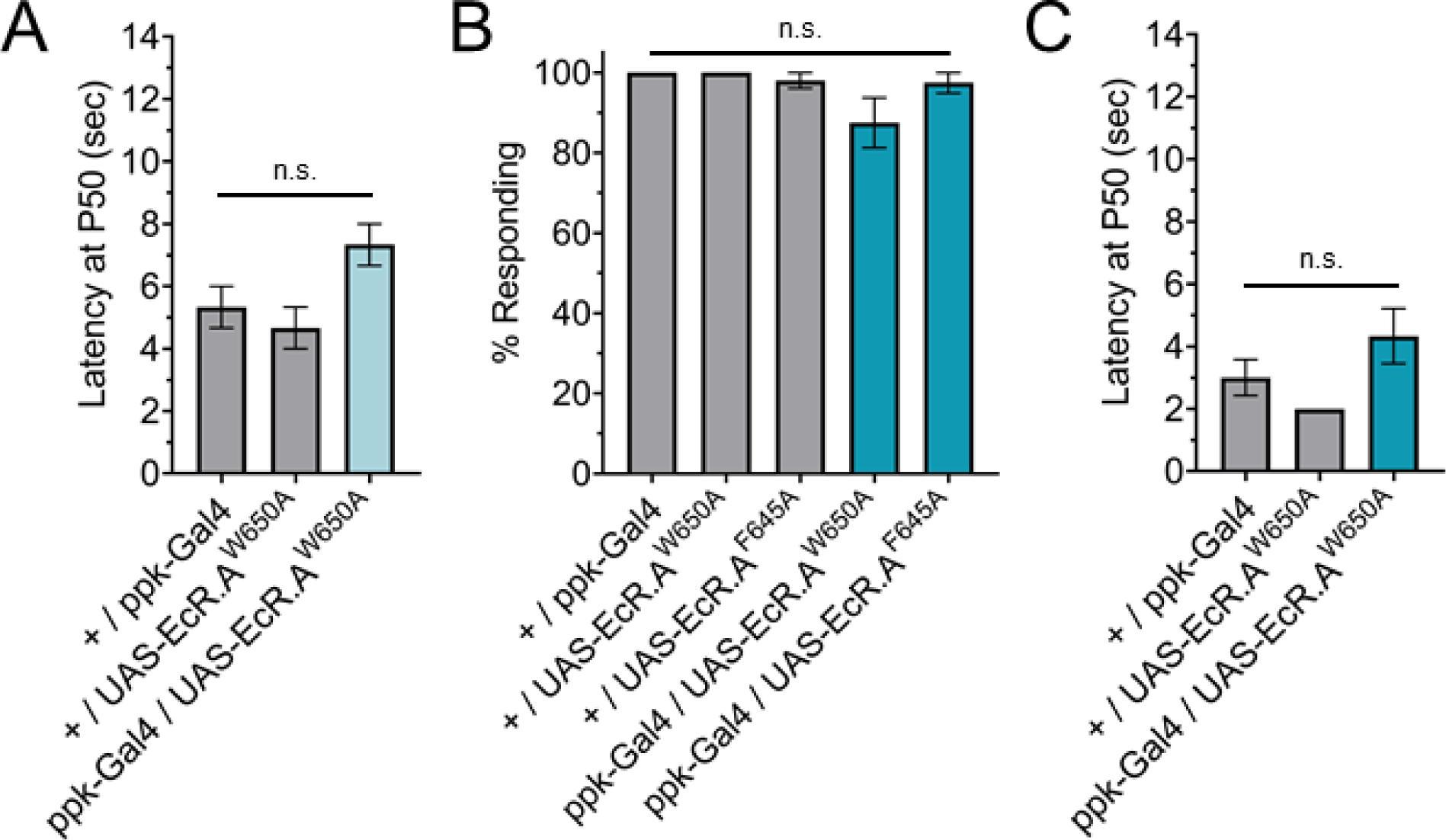
The nociceptive transition is ecdysone ligand activity dependent. (A) Latency by which 50% of the responding population displaying nociceptive behavior when overexpressing ligand binding mutant EcR-A (ppk-Gal4/UAS-EcR-A -W650A). 3^rd^ instar larvae with 42°C nociceptive probe. (B) Percent of population responding and (C) latency of 3^rd^ instar larvae with 46°C nociceptive probe. One-way Anova. n = 2-4 staging replicates of 15-20 larvae were tested for each genotype and temperature.

**Figure 4 – figure supplement 1:**
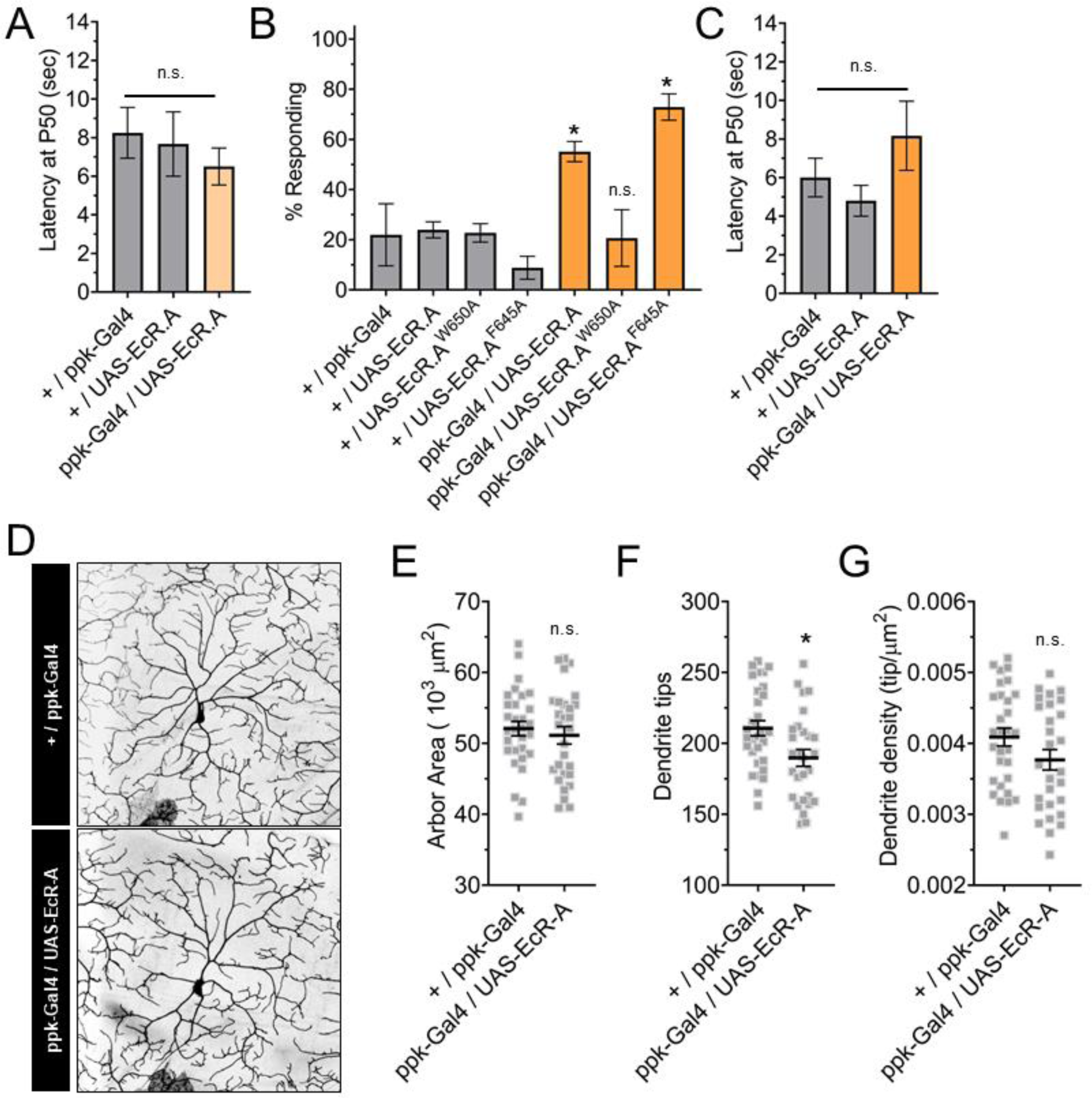
EcR-A overexpression in C4da neurons increases nociception in the 2^nd^ instar. (A) 2^nd^ instar latency by which 50% of the responding population displaying nociceptive behavior when overexpressing wildtype EcR-A. (B) Nociceptive behavior in 2^nd^ instar larvae with overexpression of EcR-A mutants. Ligand binding mutant EcR-A (ppk-Gal4/UAS-EcR-W650A), or co-activator mutant EcR-A (ppk-Gal4/UAS-EcR-F645A) with 46°C nociceptive probe. (C) Latency by which 50% of the responding population displaying nociceptive behavior. (D-F) 2^nd^ instar C4da neuron arbors overexpressing wildtype EcR-A. (D) Representative images of 2^nd^ instar C4da neuron arbors. (E) 2^nd^ instar C4da neuron arbor area. (F) 2^nd^ instar C4da neuron dendrite tip number. (G) 2^nd^ instar C4da neuron dendrite density. A-C: One-way Anova with Tukey’s. E-F: Student’s t-test. * p < 0.05. n = 2-4 staging replicates of 15-20 larvae were tested for each genotype.

**Figure 4 – figure supplement 2:**
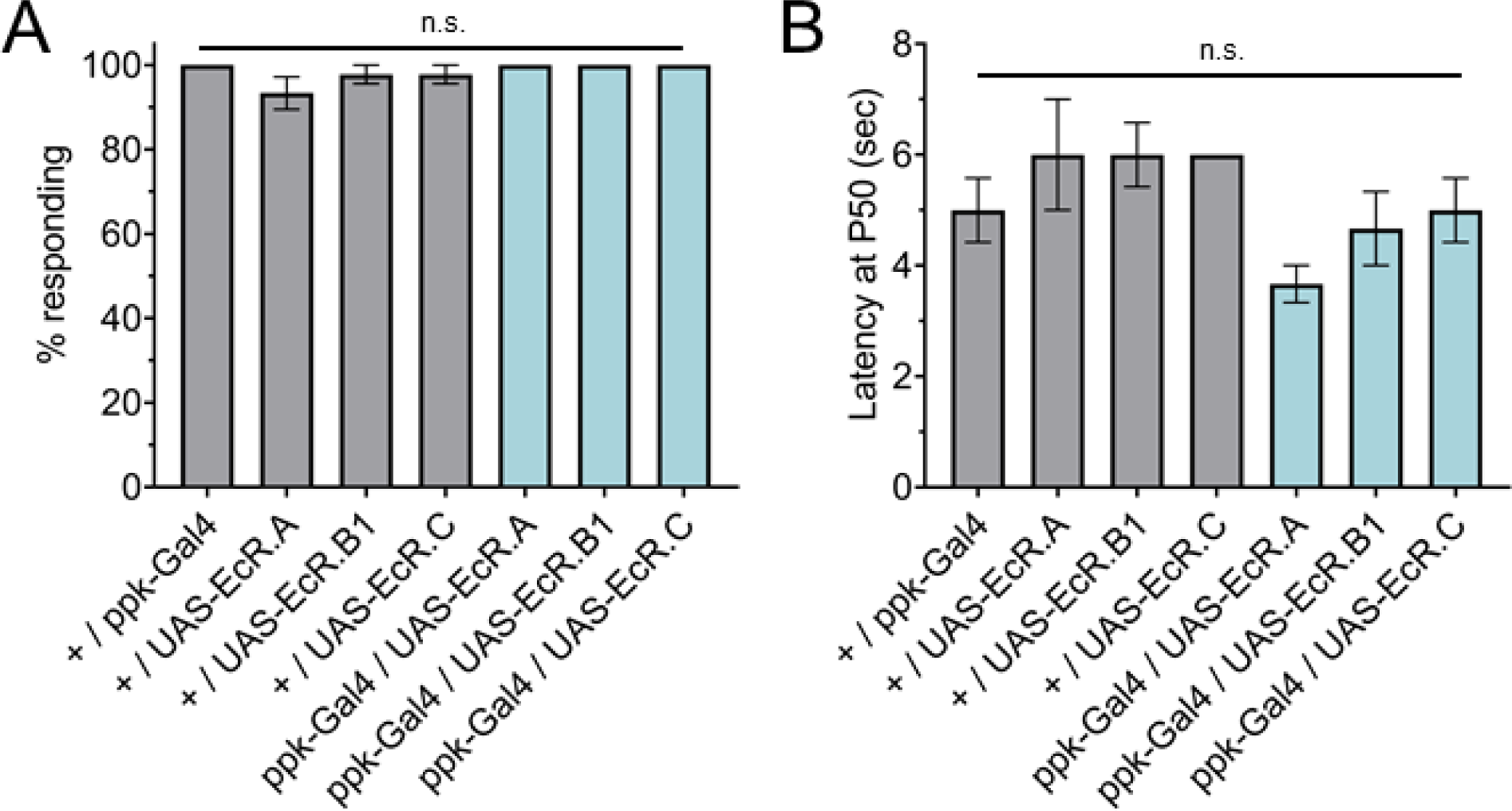
EcR isoform overexpression in C4da neurons does not change nociception in the 3^rd^ instar. (A) Nociceptive behavior in 3^rd^ instar larvae with EcR isoforms overexpressed in C4da neurons. (B) 3^rd^ instar latency by which 50% of the responding population displaying nociceptive behavior when overexpressing EcR isoforms. 42°C nociceptive probe. One-way Anova with Tukey’s. n = 3 staging replicates of 15-20 larvae were tested for each genotype.

**Figure 5 – figure supplement 1:**
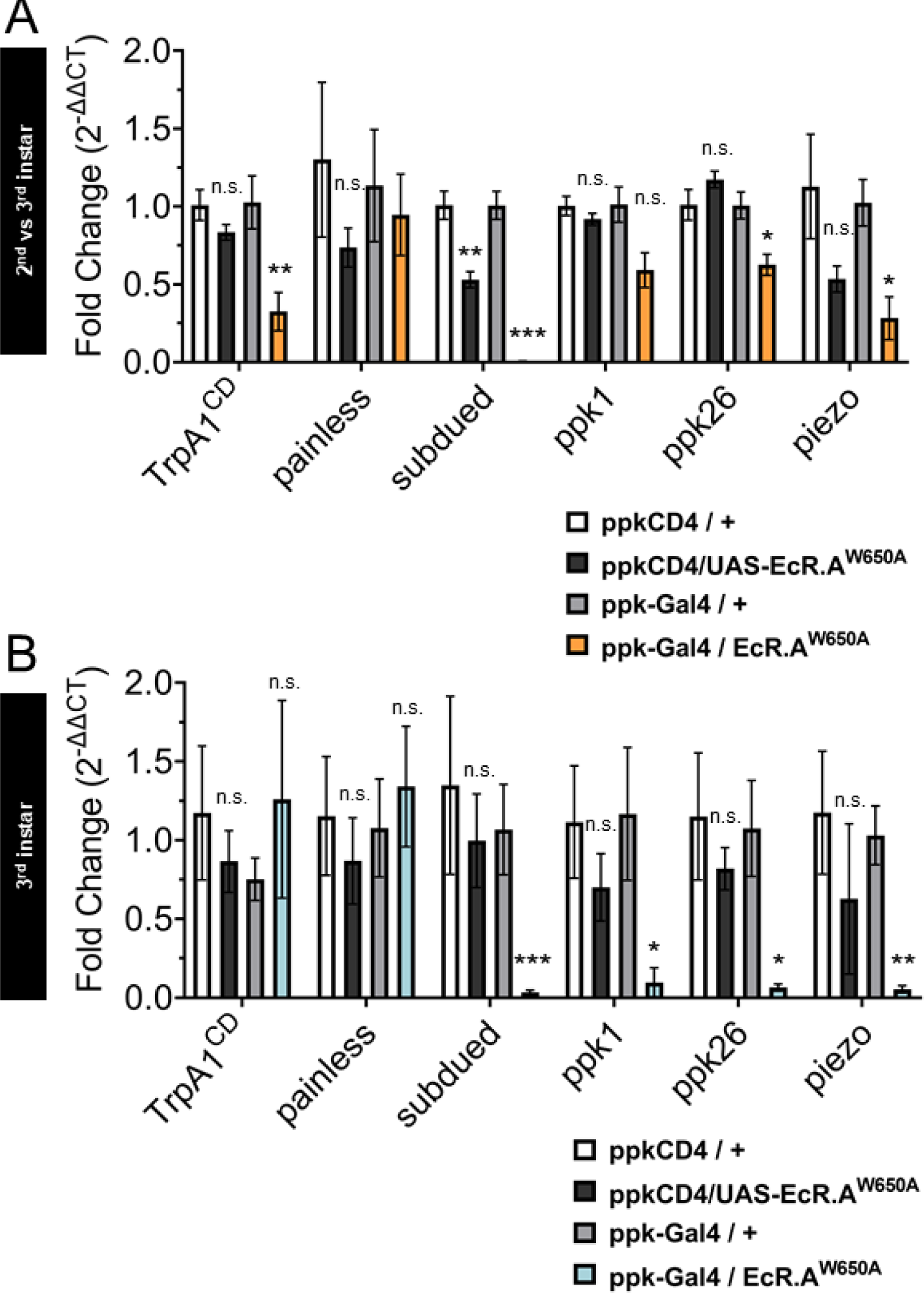
Mutant Ecdysone Receptor transcriptionally regulates nociceptive genes during the period of thermal nociceptive development. Expression of nociceptive genes measured by qRT-PCR from FACS purified C4da neurons. (A) 2^nd^ instar and (B) 3^rd^ instar C4da neurons expressing ligand binding mutant EcR-A-W650A. Genotypic controls not expressing EcR-A-W650A (ppkCD4/UAS-EcR-A-W650A) were normalized to age matched neurons (ppkCD4/+). Expression in EcR-A-W650A expressing neurons (ppk-Gal4/UAS-EcR-A-W650A) was normalized to control neurons without EcR-A-W650A expression (ppk-Gal4/+). Student’s t-test. * p < 0.05, ** p < 0.01, *** p < 0.001. n = 3-4 FACS isolation/staging replicates for each genotype.

**Figure 5 – figure supplement 2:**
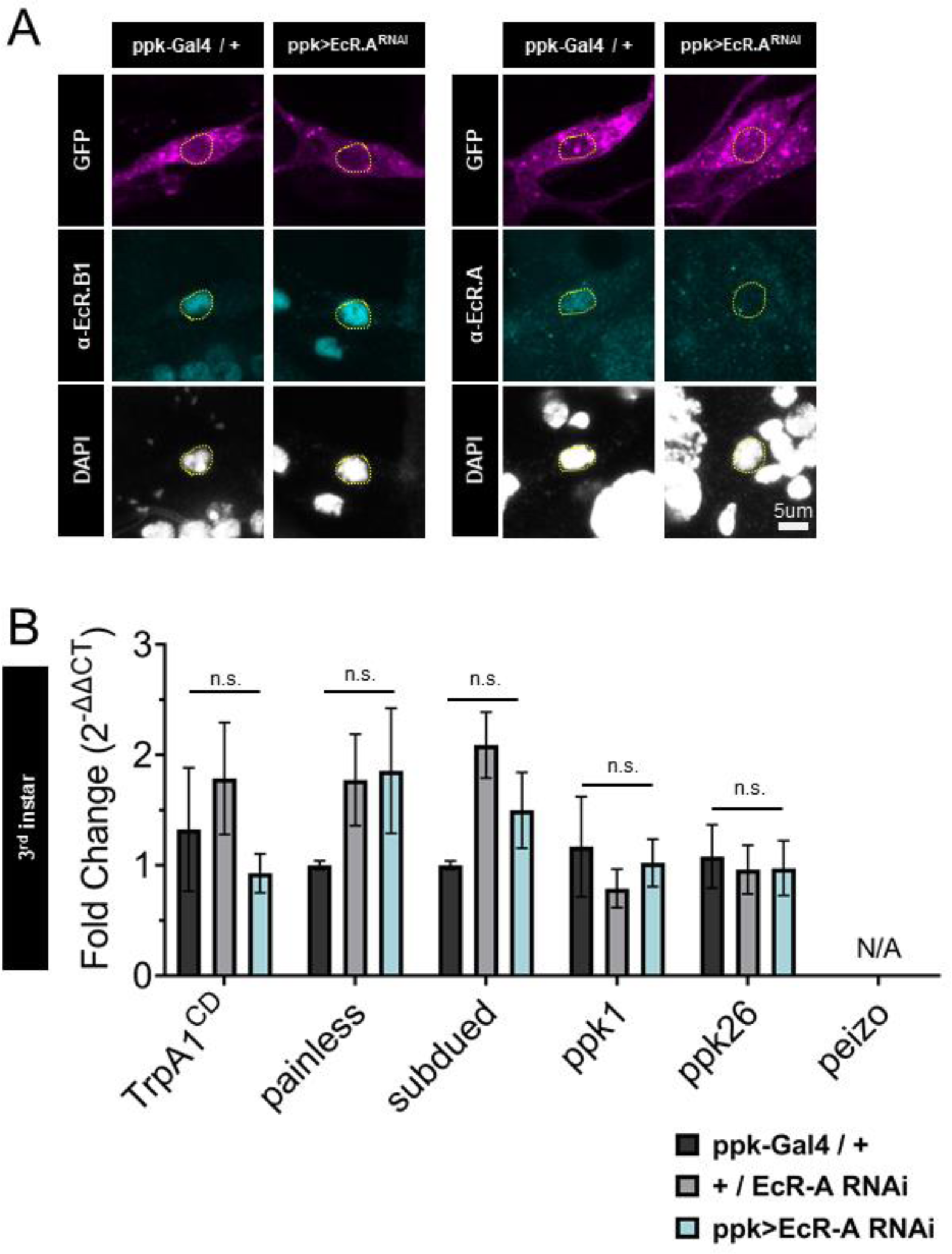
Ecdysone Receptor A (EcR-A) RNAi reduces EcR-A protein but does not alter nociceptive gene expression. (A) EcR-A RNAi reduces EcR-A protein expression and does not reduce EcR-B1 protein expression. EcR immunohistochemistry with EcR-B1 or EcRA antibody. Yellow dashed line indicates position of nucleus. C4da neurons labeled by ppktd-GFP. (B) C4da neurons isolated from 3^rd^ instar larvae expressing EcR-A RNAi. Genotypic controls not expressing EcR-A RNAi (+/EcR-A RNAi) and EcR-A RNAi expressing neurons (ppk>EcR-A RNAi) were normalized to age matched control neurons without EcR-A RNAi expression (ppk-Gal4/+). Transcripts of *piezo* were not measurable from samples (N/A: not applicable). Student’s t-test. n = 3-4 FACS isolation/staging replicates for each genotype.

**Figure 6 – figure supplement 1:**
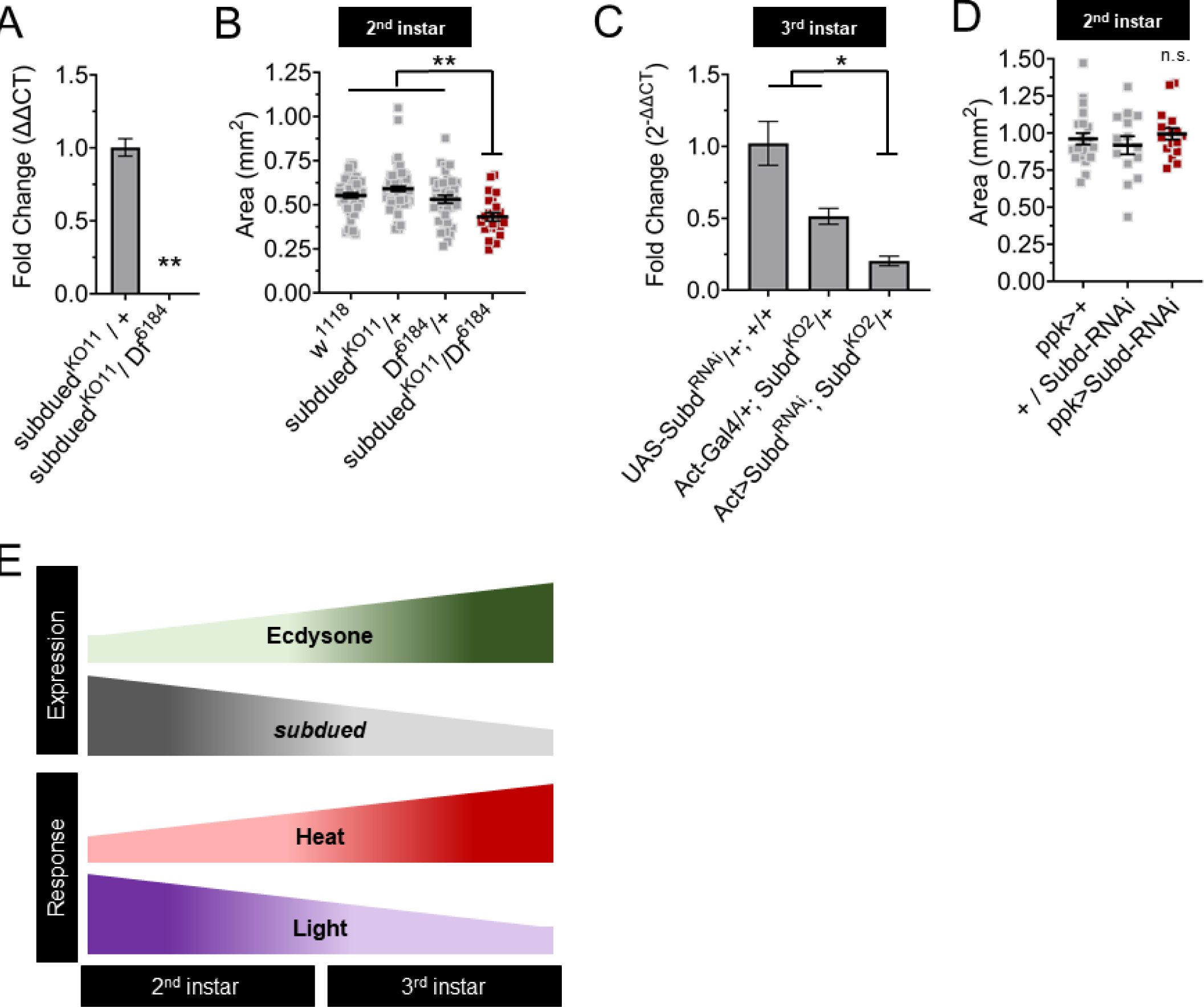
Subdued mutants have reduced larvae growth during the 2^nd^ instar. (A) *subdued* expression in knockout mutant larvae measured by qRT-PCR. (B) Size of 2^nd^ instar larvae mutant for subdued measured by body area. (A) *subdued* expression in Subdued RNAi expressing larvae measured by qRT-PCR. (D) Size of larvae expressing subdued-RNAi in C4da neurons. (E) Ecdysone regulates *subdued* expression to regulate the sC4da neuron sensory switch. A: Student’s t-test. B-D: One-way Anova. * p < 0.05. ** p < 0.01. A,C: n = 3 whole larvae isolation replicates.

